# Melanoma clonal subline analysis uncovers heterogeneity-driven immunotherapy resistance mechanisms

**DOI:** 10.1101/2023.04.03.535074

**Authors:** Charli Gruen, Howard H. Yang, Antonella Sassano, Emily Wu, Vishaka Gopalan, Kerrie L. Marie, Andrea Castro, Farid Rashidi Mehrabadi, Chih Hao Wu, Isabella Church, Gabriel A. Needle, Cari Smith, Sung Chin, Jessica Ebersole, Christina Marcelus, Anyen Fon, Huaitian Liu, Salem Malikic, Cenk Sahinalp, Hanna Carter, Sridhar Hannenhalli, Chi-Ping Day, Maxwell P. Lee, Glenn Merlino, Eva Pérez-Guijarro

## Abstract

Intratumoral heterogeneity (ITH) can promote cancer progression and treatment failure, but the complexity of the regulatory programs and contextual factors involved complicates its study. To understand the specific contribution of ITH to immune checkpoint blockade (ICB) response, we generated single cell-derived clonal sublines from an ICB-sensitive and genetically and phenotypically heterogeneous mouse melanoma model, M4. Genomic and single cell transcriptomic analyses uncovered the diversity of the sublines and evidenced their plasticity. Moreover, a wide range of tumor growth kinetics were observed *in vivo*, in part associated with mutational profiles and dependent on T cell-response. Further inquiry into melanoma differentiation states and tumor microenvironment (TME) subtypes of untreated tumors from the clonal sublines demonstrated correlations between highly inflamed and differentiated phenotypes with the response to anti-CTLA-4 treatment. Our results demonstrate that M4 sublines generate intratumoral heterogeneity at both levels of intrinsic differentiation status and extrinsic TME profiles, thereby impacting tumor evolution during therapeutic treatment. These clonal sublines proved to be a valuable resource to study the complex determinants of response to ICB, and specifically the role of melanoma plasticity in immune evasion mechanisms.

## INTRODUCTION

Somatic mutations accumulated during cancer progression and cellular plasticity driven by epigenetic and transcriptomic regulation confer to solid tumors high complexity and adaptability to (micro)environmental stressors^1,2^. Consequently, intratumor heterogeneity (ITH) has been identified as a key element contributing to metastatic potential and therapeutic resistance^3–6^. Moreover, the changes that occur over space and time in the tumor microenvironment (TME) shape the intratumoral landscape, thereby guiding the subclonal evolution of cancer cells^7–9^. The impact of ITH on conventional and targeted therapy is well established3. For example, most patients with metastatic melanoma who responded to BRAF and MEK inhibitors eventually relapsed due to subpopulations of cells with distinct phenotypes and variability in expressed mutations10. However, the mechanisms governing ITH and its specific functions are not fully understood, especially those related to immunotherapy response^6,11^.

Immune checkpoint blockade (ICB) is currently the first-line treatment for unresectable or metastatic melanoma; however, disease resistance and relapse remain a clinical issue. Correlates of clinical benefit include high tumor mutational burden (TMB), immunogenic neoantigen load, and increased tumor-infiltrating lymphocytes, as ICB efficacy relies on the anti-tumor activity of T cells in the tumor microenvironment (TME)^12^. In contrast, defects in antigen presentation and Interferon gamma (IFNγ) response are associated with poor outcomes^11^. However, the discovery of definitive biomarkers of ICB efficacy that allow patient stratification is still an urgent unmet need. We previously developed a panel of four (M1-M4) genetically engineered mouse (GEM) models of cutaneous melanoma and performed comparative analyses of their genomic, transcriptomic, and immune cell profiles^13^. The results demonstrated that M1-M4 are representative of multiple molecular and phenotypic subtypes of human melanomas and comprise a preclinical platform to study molecular mechanisms of immunotherapies. Importantly, this study identified a mouse-derived signature, the melanocytic plasticity signature (MPS), predictive of patient outcomes in response to anti-CTLA-4 or anti-PD-1, which linked, for the first time, melanoma multipotency and undifferentiation profiles with ICB resistance.

A range of strategies exist to investigate intratumoral clonal dynamics based on barcoding, computational tracing, and spatial molecular profiling; however, they are limited in their ability to dissect genetic versus non-genetic determinants of tumor progression and therapeutic response. Here we address this challenge with the generation of a novel model system comprised of 24 single cell-derived clonal sublines derived from M4 melanoma. We characterized the 24 clones with genomic and single cell transcriptomic analyses accompanied by tumor growth *in vivo* and anti-CTLA-4 efficacy studies. Heterogeneity within sublines was observed for both phenotypic subtypes and TME profiles, linking heterogeneity with tumor behavior and providing further insight into the role of melanoma plasticity in immune evasion mechanisms.

## RESULTS

### Generation of 24 single-cell derived clonal sublines from ICB-sensitive mouse melanoma

To build a model system for dissecting genetic and non-genetic factors that contribute to intratumoral heterogeneity in response to ICB, we established 24 single cell-derived clonal sublines (C1-C24) from a genetically engineered mouse melanoma (M4)^13^. M4 was developed by exposing hepatocyte growth factor (HGF) transgenic mice to neonatal ultraviolet (UV) radiation^14^ and exhibits high mutational burden, histological and phenotypic heterogeneity, and a transitory developmental phenotype^13,15^. Notably, M4 is highly sensitive to ICB, reaching 40-60% response rates that range from full regression to significant delays in tumor growth and relapse after treatment. We sorted live single cells from M4 cell line by fluorescence-activated cell sorting (FACS) into a 96 well plate, expanded 24 sublines in culture, and generated a biobank of low passage sublines for molecular characterization and functional analyses **(Fig. 1a)**.

**Fig. 1.**
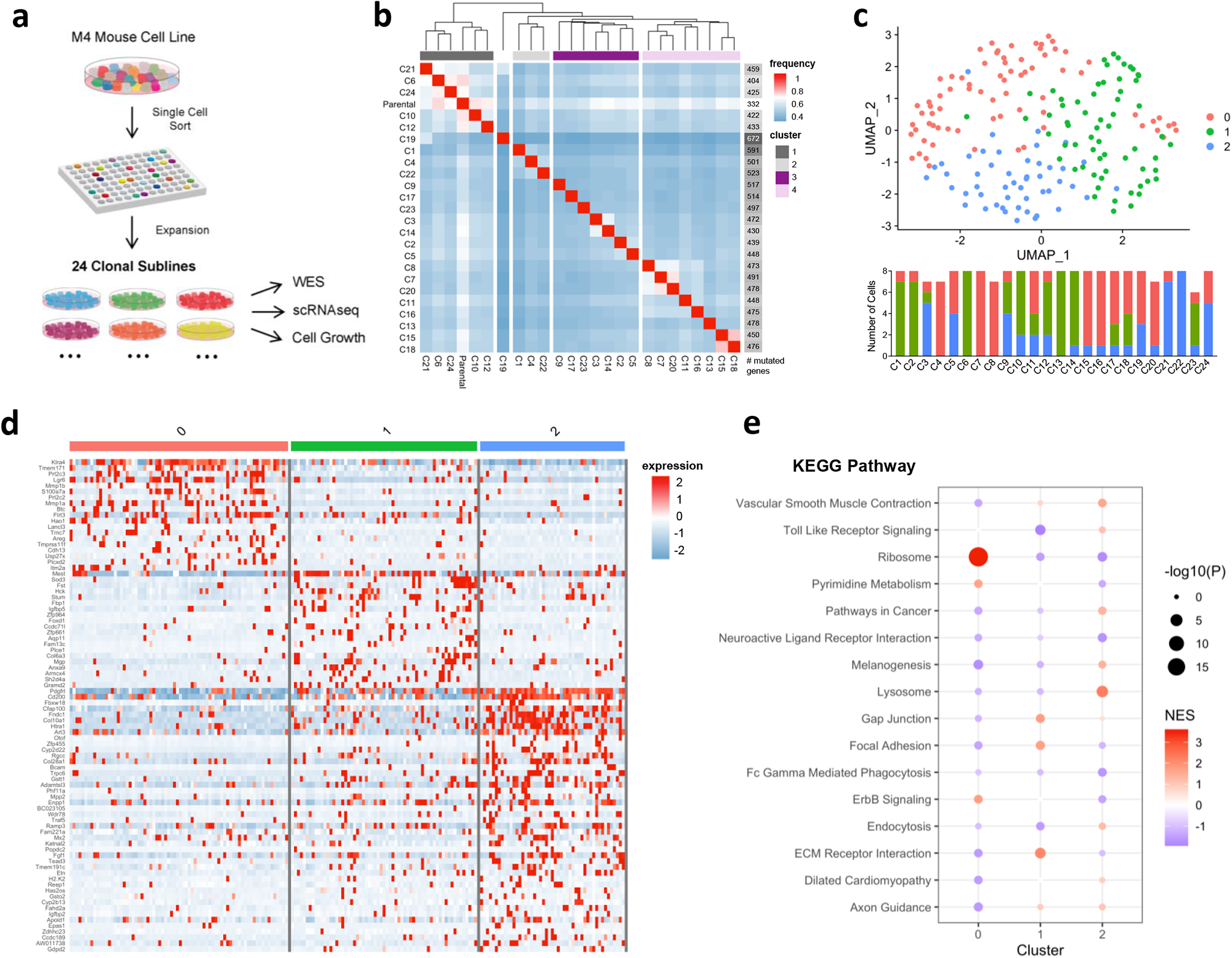
Generation of 24 single-cell derived clonal sublines from M4 mouse melanoma cell line. **a,** Individual cells from the M4 (B2905) mouse cell line were isolated for cell culture via single-cell sorting. A total of 24 single-cell derived cultures were successfully expanded to create 24 clonal sublines. Whole exome sequencing (WES), Smart-seq2 full-length single-cell RNA sequencing (scRNAseq), and IncuCyte live cell growth analysis was performed. **b,** Heat map of overlapping mutated genes detected in the clonal sublines by WES. The total number of mutated genes in each clone is indicated in the right. **c,** Single cell expression profiles obtained from scRNAseq clustered and visualized with UMAP analysis. Number of cells per cluster in each clonal subline is shown below UMAP. **d,** Heat map of top 25% differentially expressed genes per UMAP cluster based on Log_2_FC. **e,** Gene set enrichment analysis (GSEA) of KEGG pathways in each UMAP cluster. Pathways with significant (*P*≤0.05) positive or negative enrichment in at least one UMAP cluster (0-2) are shown. NES: normalized enrichment score.

To characterize the mutational landscape of the 24 clones, whole-exome sequencing (WES) analysis was performed on the cultured sublines. Genetic diversity was revealed by nonsynonymous single nucleotide variant (SNV) calling. Among the unique mutations distinguishing the clones, we identified genes related to hypoxia (*Hif1a*, C1), inflammatory response (*Cd8a*, C11; *Il22* and *Il10ra*, C1; *Il17re*, C14; *Map3k7*, C15), chromatin remodeling and replication (*Atrx*, C11), and drug resistance (*Abcc5*, C18) (**Table S1**). Additionally, we analyzed the frequency of overlapping mutated genes among the 24 clonal sublines and the parental M4 cell line and identified four main branching groups, illustrating the genetic complexity of M4 melanoma **(Fig. 1b)**. The branching groups were consistent with the evolutionary relationships inferred from *Trisicell*^16^, a computational toolkit we developed for the phylogenetic analysis of tumors based on SNVs at the single cell level **(Fig. S1a)**. Moreover, the phylogeny tree built using the expressed SNVs called from single-cell RNA sequencing^17^ (scRNAseq) data of the clonal sublines grown in culture, confirmed the genetic intra-clonal homogeneity of the cells from each clone **(Fig. S1b)**.

For an unbiased characterization of the transcriptomes of the 24 clonal sublines, we performed UMAP (Uniform Manifold Approximation and Projection) analysis of the scRNAseq data, and three clusters (0-2) of cells were distinguished **(Fig. 1c)**. Although approximately 50% of the clones were fully composed of cells mapped in a single UMAP cluster, multiple clonal sublines contained cells exhibiting 2 or even 3 distinct transcriptomic profiles (e.g., C11, C18, C23, C12, C9 and C17), suggesting elevated plasticity of these clones, **(Fig. 1c and S1c-d)**. To understand the identity of the cells in each cluster, we performed differential gene expression analysis followed by Gene Set Enrichment Analysis (GSEA) using Kyoto Encyclopedia of Genes and Genomes (KEGG) pathways **(Fig. 1d-e and Table S2-3)**. Cells in Cluster 0 were marked by an upregulation of pathways related to proliferation (e.g., ‘Ribosome’, ‘Pyrimidine metabolism’ and ‘ErbB signaling’). Cluster 1 cells were enriched in pathways related to extracellular matrix (ECM) dynamics and cell migration (e.g. ‘Focal adhesion’), which could reflect an invasive phenotype. In contrast, the pathways upregulated in Cluster 2 cells were related to melanogenesis, protein degradation and recycling (e.g., ‘Lysosome’ and ‘Endocytosis’), indicative of a more differentiated phenotype **(Fig. 1e)**.

We further investigated the expression of canonical markers of melanoma differentiation states (e.g., *Mitf, Dct, Ngfr, So×10, Erbb3, Axl*)^15^ and the melanocytic plasticity signature (MPS), which was correlated with response to ICB therapy^13^ **(Fig.S2)**. Overall, most of the cells showed elevated MPS scores regardless of the clonal subline, indicative of undifferentiated or neural crest-like states and, therefore, the UMAP clusters did not fully separate clones with distinct developmental states **(Fig.S2a-c)**. However, some developmental markers were significantly enriched in specific clusters. For example, *Axl* and *Ngfr* expression were increased in cluster 0 while *So×10* and *ErbB3* levels were the highest in cluster 2, which could reflect a transition phenotype towards a more differentiated state **(Fig.S2d)**. Altogether, these results demonstrate the marked diversity of the 24 clones at both genetic and transcriptional levels, uncovering melanoma cell intrinsic pathways that could have implications for their potential to survive and form tumors *in vivo^18^*.

### The melanoma clonal sublines exhibit a wide range of tumor growth kinetics

To evaluate the tumorigenic potential of the 24 clones, we implanted 1 million cells of each subline into syngeneic C57BL/6 mice and measured tumor size until endpoint, which defined the survival time (see **Methods**). We observed a broad spectrum of tumor growth kinetics, resulting in a range of survival outcomes falling above and below the survival of parental M4 melanoma **(Fig. 2a-b).** While most of the clonal sublines (16 clones) formed tumors in 100% of the mice (N=5/5), only 60% of mice implanted with clones C7, C10 and C20 (N=3/5) and 40% with clones C5 and C21 (N=2/5) reached endpoint within 7 months. Clones C6, C13 and C19 were not capable of forming tumors in immunocompetent mice (N=0/5). The clonal sublines that formed 3 or more tumors were grouped by quartiles of median survival (G1-G4) and those not reaching median survival by the end of the study were assigned to group 5 (G5; **Fig. 2a-b)**. Of note, only 2 clonal sublines (C15 and C18) showed faster tumor growth than the parental M4, whereas all mice implanted with clones in G2-G5 presented significantly improved survival (**Fig. 2b**). These results illustrate the phenotypic diversity of the 24 clonal sublines *in vivo* and suggest that some extent of genetic ITH is an advantage for tumor fitness, in line with previous findings^19^.

**Fig. 2.**
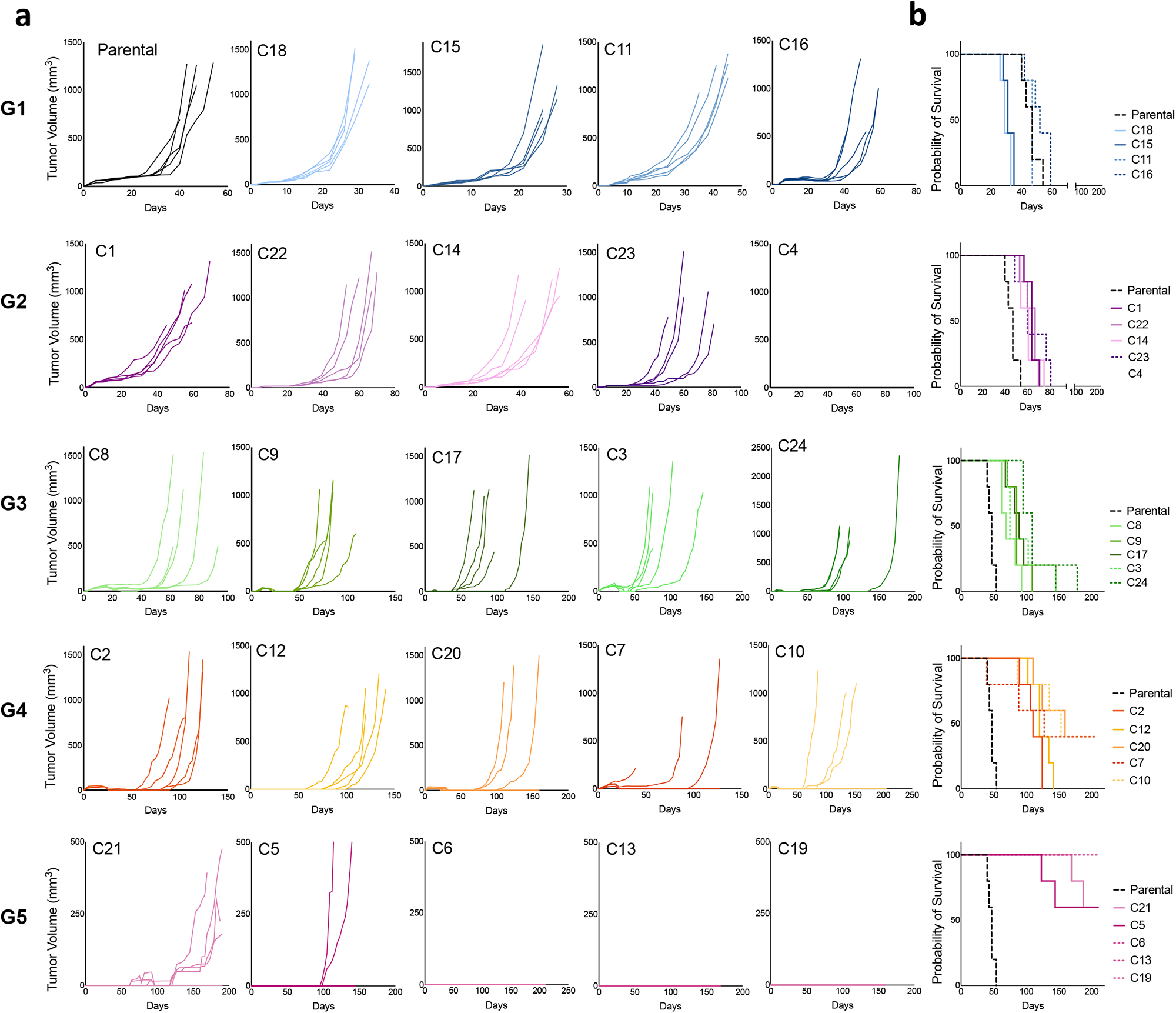
Diverse tumor growth kinetics of the 24 clones in syngeneic mice. 1.0 x 10^6^ melanoma cells from the 24 clonal sublines were implanted subcutaneously into C57BL/6 syngeneic mice. **a,** Tumor growth curves of the clones implanted in untreated mice (N=5 mice per clone). Median survival time grouped by quartiles (Q) was used to establish five survival groups (G1-G5), each labeled with a different color: G1<Q1 (blue), G2=Q1-median (purple), G3=median-Q3 (green), G4>Q3 (orange), G5<3 tumors formed (pink). **b,** Kaplan-Meier survival curves from Fig.2a. Parental M4 is included for reference in each group.

To address whether the observed tumor growth curves were determined by the proliferative ability of the 24 clones, we analyzed cell growth kinetics in culture using IncuCyte live imaging **(Fig. 3a)**. Cells from each of the clones were plated at similar densities and their confluency was estimated every 3 hours for 5 days. Most of the curves followed a sigmoid shape and plateau between 50 to 120 hours at 85-99% confluency. Doubling times ranged from 17 to 33 hours, with C18, C1 and C15 being the fastest growers while C11, C13, and C19 were the slowest. When all survival groups were analyzed together, we found a significant correlation between median survival *in vivo* and doubling times *in culture*, with C5, C6 and C11 being clear outliers (**Fig. 3b**). In line with this observation, the clonal sublines with the fastest tumor growth (low median survival) tended to map in UMAP Cluster 0, which was enriched in proliferation-related pathways (e.g., ‘Ribosome’ and ‘Pyrimidine metabolism’), although no significant differences between clusters were found (**Fig. 3c and 1e**). Together, these results indicate that cell proliferation kinetics contribute to the tumor growth of the 24 clonal sublines *in vivo*, although additional mechanisms may explain their tumorigenicity.

**Fig. 3.**
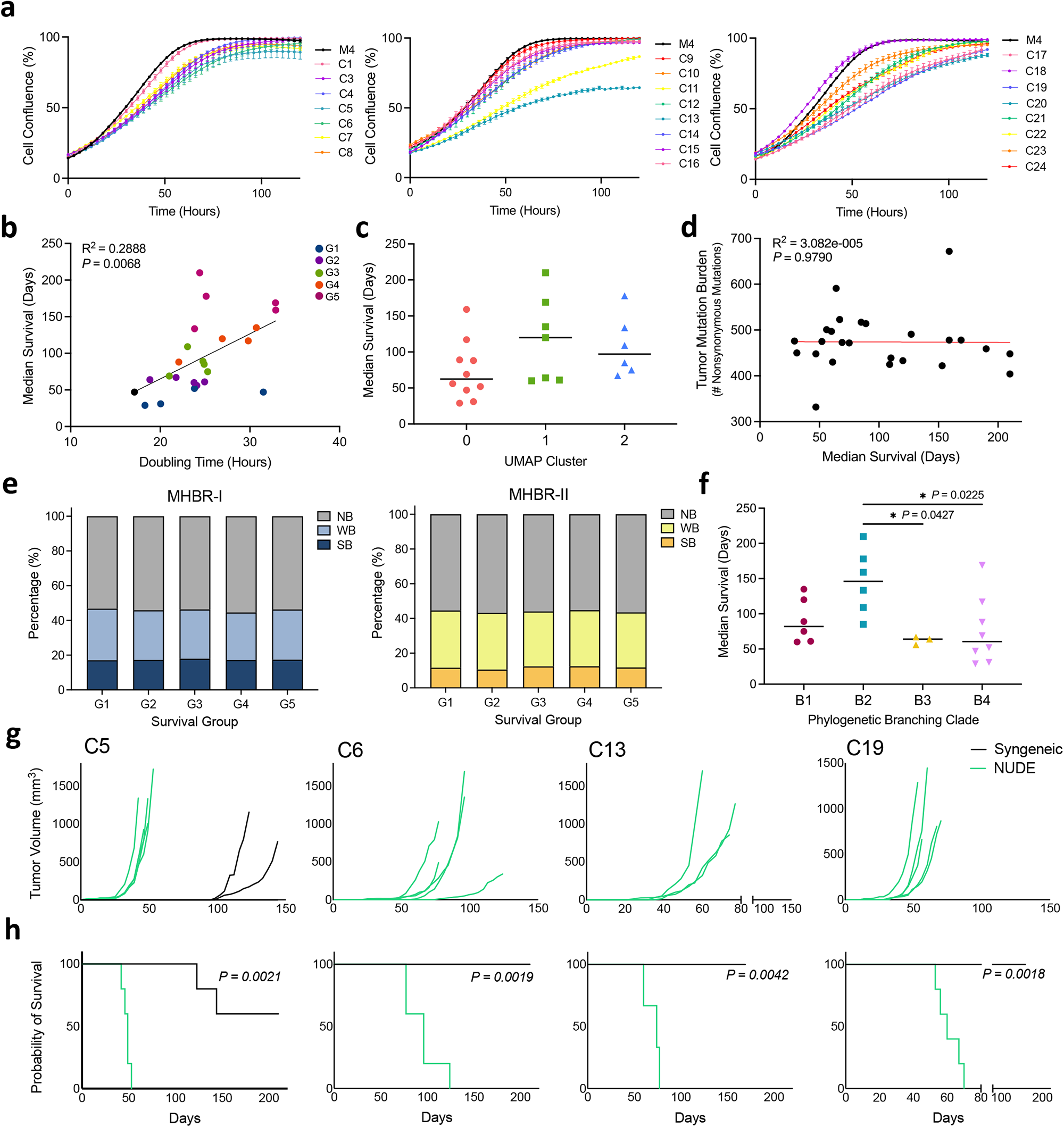
Correlation between cell growth, mutational profiles and the tumorigenicity of the clonal sublines. **a,** Cell growth kinetics of the 24 clones. Cell confluency was estimated for 5 days using IncuCyte live imaging analysis. **b,** Correlation of the median survival of mice implanted with the 24 clonal sublines *in vivo* and the doubling time of the 24 clonal sublines *in culture*. Colors correspond to survival groups from Fig.2. Pearson coefficient (R^2^) and *P* value are indicated. **c,** Median survival of the clones in each UMAP cluster from Fig.1c. No significant difference between clusters were found by one-way ANOVA test adjusted for multiple comparisons. **d,** Correlation analysis of the median survival of the mice implanted with the 24 clonal sublines and the tumor mutation burden of each clone determined by WES. Pearson coefficient (R^2^) and *P* value are indicated. **e,** MHBR-I and MHBR-II binding prediction of mutated genes from WES data. Binding scores were calculated individually per clone and the percentage of mutations falling into three categories (NB: non-binding, WB: weak-binding, SB: strong-binding) is represented. Results are shown as the mean for each survival group (G1-G5, Fig.2). **f,** Median survival of the clones and corresponding phylogenetic branches from Fig.S1a. *P* values shown for one-way ANOVA test adjusted for multiple comparisons. **g,** 1.0 x 10^6^ melanoma cells from C5, C6, C13, and C19 (group G5 in Fig.2) were implanted subcutaneously into immunocompetent syngeneic C57BL/6 (black lines, N=5) or immunodeficient NUDE (green lines, N=5) mice. **h,** Kaplan-Meier survival curves from Fig.3g. Two-tailed *P* values from log-rank (Mantel-Cox) test are indicated.

### Poor tumor formation capacity is associated with increased immunogenicity of the clonal sublines

High TMB in melanoma resulting in the expression of neoantigens promotes anti-tumor immune responses^20,21^. To investigate the potential correlation between TMB and tumor growth of the clones, we compared the number of nonsynonymous mutations from WES of the clonal cells to the median survival of syngeneic mice implanted with each subline **(Fig. 3d)**. The fastest growing clones, C15 and C18, showed TMB below the mean and low numbers of unique mutations while C19 had the highest TMB and, along with group G5, had the longest survival outcomes **(Fig. 1b and 2b)**. However, the mutational burden of the clones did not correlate with their *in vivo* tumor growth kinetics when we compared the results among all clones **(Fig. 3d)**. Therefore, we assessed the potential immunogenicity of the mutations detected in each clone by calculating the Mouse Harmonic-mean Best Rank (MHBR) scores, a predictive assessment of the binding affinity of neopeptides generated by nonsynonymous SNVs to MHC class I (MHC-I) and MHC class II (MHC-II) receptors^22^ **(Fig. S3a)**. Among all clones, C1 and C19 had the greatest TMB and number of predicted MHC-I and MHC-II receptor binding neoantigens **(Fig.S3b)**. Overall, there was a greater percentage of predicted strong-binding (SB) neoantigens to MHC-I versus MHC-II, but no significant differences were observed between the survival groups (G1-G5) **(Fig. 3e)**. Next, we examined the correlation of tumor growth with the mutational profiles of the clones defined by the branches (B1-B4) of the phylogeny trees generated by *Trisicell*^11^ **(Fig. 3f and S1a).** Although we found no differences in cell doubling times of the sublines from each branch (data not shown), B2 corresponding clones showed significantly delayed median survival *in vivo* compared to those in B3 and B4. These results suggest that the specific mutational profiles of the clonal sublines rather than total neoantigen load may trigger anti-tumor immune responses in baseline conditions, determining the tumorigenicity of the clonal sublines.

We further explored if the lack of tumor formation observed for C5, C6, C13 and C19 was caused by anti-tumor T cell responses by implanting them into athymic NUDE mice **(Fig. 3g-h)**. All four clones were able to form tumors in NUDE mice with 100% penetrance, indicating that the antitumoral T cell responses in immunocompetent mice prevented *in vivo* growth of these 4 clonal sublines **(Fig. 3g)**. In line with the cell growth kinetics in culture, C5 showed the fastest tumor growth in NUDE mice (median survival=49 days vs. 96, 74 and 60 days for C6, C13 and C19, respectively) **(Fig. 3h)**. Our results so far demonstrate that both the proliferation capacity and the immunogenicity of the clonal sublines play an important role in their tumorigenicity. However, the link between immunogenicity and tumor growth dynamics cannot be explained by the neoantigen load alone.

### Inter- and intra-tumor heterogeneity of developmental pathway expression in melanomas from the clonal sublines

To investigate in more detail the molecular features of the clonal sublines *in vivo*, we selected 11 clones with the most consistent tumor growth kinetics to perform comparative transcriptomic and histological analyses. Three representative tumors from each clonal subline were subjected to bulk RNA sequencing (bRNAseq) **(Fig. 4a)**. Principal component analysis (PCA) revealed marked heterogeneity between tumors, even those from the same clonal sublines **(Fig. 4b)**. Unsupervised hierarchical clustering using the top 300 most variably expressed genes between the 33 tumors separated them into four main clades (I-IV) **(Fig. 4c)**. Notably, almost half of the clones showed inter-tumor heterogeneity as not all their tumors clustered together in the same clade (e.g., C4, C8, C12, C14, C22; **Fig. 4c**). Among the top inter-clade differentially expressed genes, we found that those related to melanogenesis and melanin biosynthesis (*Dct, Tyr, Tyrp1*), glutathione metabolism (*Gsta1, Gsta2, Gsta4*) and collagen formation (*Col26a1*) were highly expressed in Clades I and II, whereas Clade IV showed upregulation of genes related to muscle contraction and myogenesis (*Mybpc1, Mybpc2, Actn2*), Wnt signaling (*Wnt10a, Wnt7b, Actc1, Myh3*), and the neuronal system (*Kcna1, Gpr17, Grin2c*) **(Fig. 4c, S4 and Table S4)**. These results suggest that one of the major distinctions of the clones *in vivo* is their differentiation status.

**Fig. 4.**
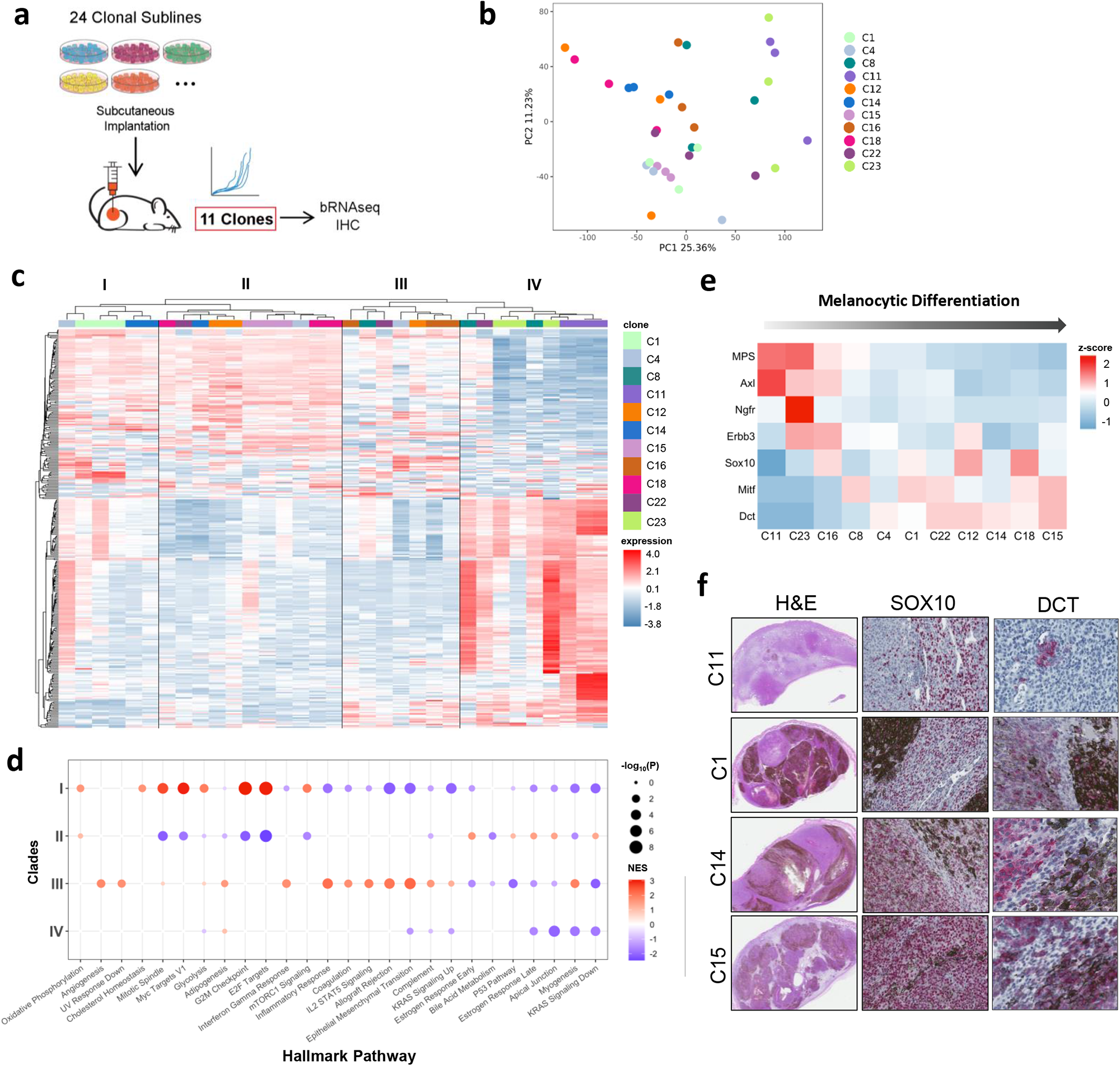
Distinct transcriptomes and differentiation status of the clones *in vivo*. **a,** After implantation into C57BL/6 mice, 11 clones with consistent and representative tumor growth kinetics were selected for molecular characterization by bulk RNA sequencing (bRNAseq) and immunohistochemistry (IHC). **b,** Principal Component Analysis (PCA) of bRNAseq data from tumors from 11 clones (N=3 tumors per clone). **c,** Heat map of the top 300 most variable genes between tumors. Unsupervised hierarchical clustering formed four distinct clades indicated as I, II, III, and IV. Colors in legend distinguish the 11 clones. Scale bar represents gene expression values after quantile normalization of raw counts. **d,** GSEA comparing the tumors in each clade from Fig.4b. Normalized enrichment score (NES) is indicated by the scale bar. **e,** Expression of melanocytic differentiation marker genes^15^ and the Melanocytic Plasticity Signature (MPS)^13^ by the clones *in vivo* (N=3 tumors per clone). **f,** Representative images of tumor sections from C11, C1 C14 and C15 stained by H&E, SOX10 or DCT (TRP2) immunohistochemistry.

GSEA revealed significant Hallmark pathways associated with the tumors in the four clades **(Fig. 4d)**. Clade I tumors showed marked upregulation of ‘Oxidative phosphorylation’, ‘Glycolysis’ and ‘Cholesterol homeostasis’ gene sets while Clades III and IV presented increased gene expression corresponding to ‘Adipogenesis’, which reflects distinct metabolic adaptation of the tumors in these clades. In line with high metabolic tumors, Clade I tumors also had enriched expression of gene sets related to proliferation, such as ‘mTORC1 signaling’, ‘Myc targets’, ‘E2F targets’, ‘G2M checkpoint’, and ‘Mitotic spindle pathway’. In contrast, Clade II showed downregulation of the main proliferation pathways and upregulation of ‘Estrogen response’. Clade III tumors exhibited enrichment of pathways related with migration and metastasis, such as ‘Myogenesis’, ‘Epithelial to mesenchymal transition’ (EMT) and ‘Angiogenesis’, which were downregulated in Clade IV. In addition, Clade III showed elevated inflammatory pathways, including ‘Inflammatory response’, ‘Interferon gamma response’, ‘IL2-STAT5 signaling’, ‘Complement’ and ‘Allograft rejection’, all of which decreased in Clade I **(Fig. 4d)**. Notably, gene expression in the tumors differed greatly from the previously described gene expression of the cells in culture, indicating the occurrence of transcriptional changes as the cells adapted to TME conditions.

To further characterize the developmental states of the 11 clonal sublines *in vivo*, we calculated their MPS scores. Consistent with previous observations, high MPS scores were correlated with more undifferentiated and neural crest-like phenotypes (C11, C23, C16) while lower MPS corresponded to transitory (C8, C4, C1, C22) and melanocytic (C12, C14, C18, C15) states, as defined by the canonical markers *Axl, Ngfr, Erbb3, Sox10, Mitf* and *Dct* **(Fig. 4e)**. We validated these findings in tumor sections stained with SOX10 or DCT antibodies **(Fig. 4f)**. Tumors with more undifferentiated phenotypes (C11, C23, C16) had little to no melanin pigmentation, lower expression of SOX10 and non-detectable levels of DCT, while tumors falling into transitory and more differentiated phenotypes (C1, C12, C14, C15) had moderate to high levels of melanin pigmentation and higher expression of SOX10 and DCT **(Fig. 4f)**. These results highlight a singular diversity of developmental states exhibited by the clonal sublines *in vivo*. Recent studies have correlated melanoma differentiation with response to immunotherapy^13,23^, underlining the value of these clones as preclinical models for immunotherapeutic treatment studies.

### Distinct tumor microenvironment composition of the melanomas from the clonal sublines

High intratumoral T cell infiltration is a hallmark of more immunogenic or “hot” melanomas with increased likelihood to respond to immunotherapy^24^. To examine T cell infiltration in the tumors from the 11 clones, we performed CD3 immunohistochemistry (IHC) and quantified CD3^+^ cells using HALO Image Analysis software **(Fig. 5a and S5)**. To evaluate whether melanin content could interfere with CD3 staining detection, a Random Forest machine learning algorithm was used to classify the regions of the tumors based on melanin levels **(Fig.S5a;** see also Methods section). We did not observe obvious differences between high and low melanin regions **(Fig.S5b-c)**; therefore, we estimated T cell infiltration using the sum of CD3^+^ cells in whole sections. C1 and C8 tumors showed the highest levels of T cell infiltration, which was comparable with parental M4 tumors. In contrast, C4, C12 and C14 exhibited the lowest amount of intratumoral CD3^+^ cells **(Fig. 5a)**.

**Fig. 5.**
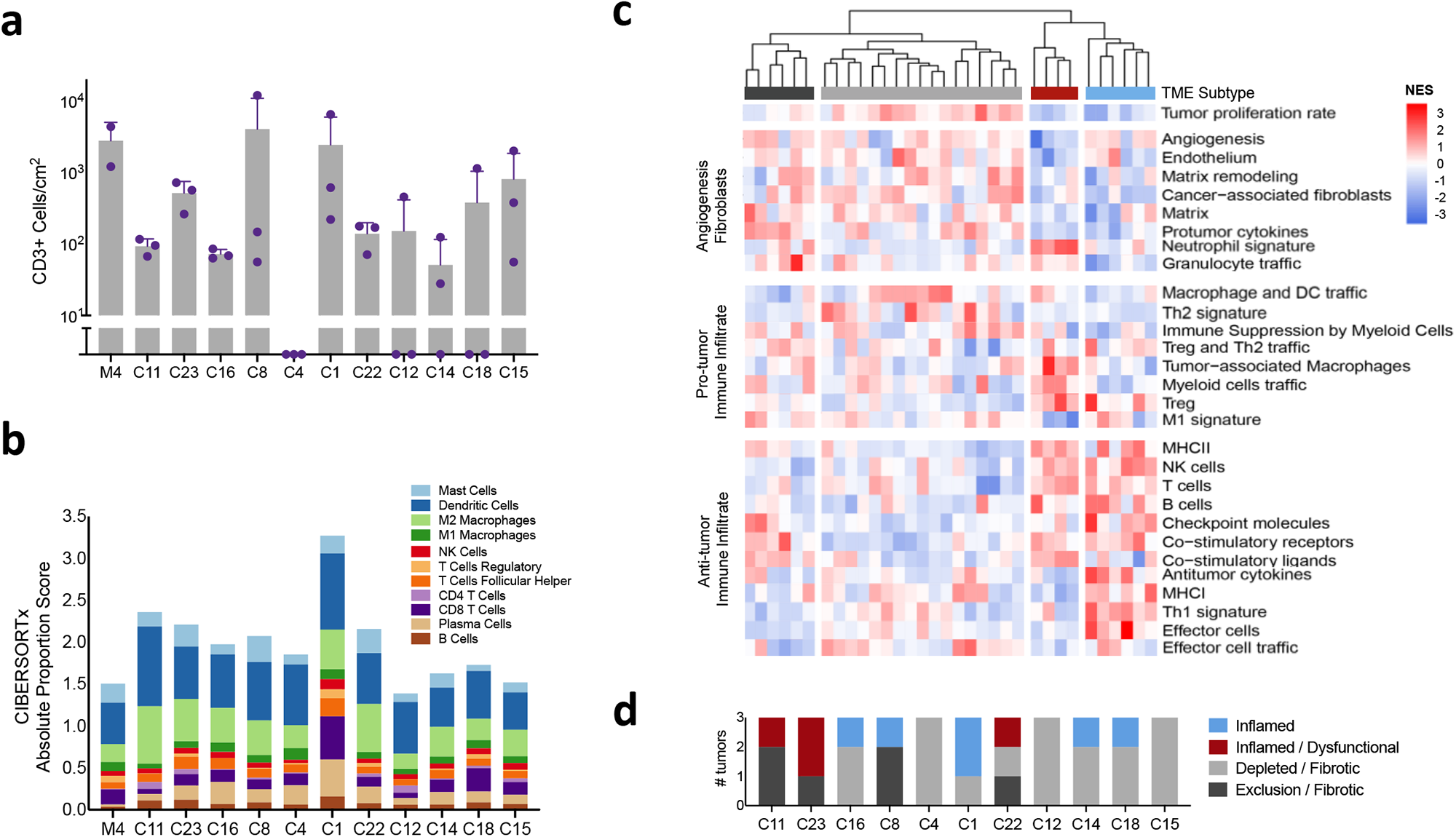
Characterization of the tumor microenvironment (TME) of the clones. **a,** CD3^+^ cells infiltrating tumors from the clones. CD3 immunostaining was quantified on whole-tumor sections using HALO Image Analysis Platform (N=3 per clone, N=2 for parental M4). **b,** CIBERSORTx analysis was performed to estimate the absolute proportions of immune cell populations in untreated tumors (N=3 per clone, N=4 for parental M4). Bar graph illustrates the average proportion of each immune cell type in tumors from each clonal subline and parental M4. Immune cells with proportions less than 0.1 are not shown. **c,** Single-sample GSEA (ssGSEA) was performed using 29 immune-related signatures^26^ on the bulk RNA sequencing data from untreated tumors. Hierarchical clustering separated the tumors into four TME groups, indicated by the color bars (dark gray: exclusion-fibrotic, light gray: depleted-fibrotic, red: inflamed-dysfunctional, blue: inflamed). Scale bar indicates range of normalized enrichment scores (NES). **d,** Classification of tumors into the four immune microenvironment categories from Fig. 5c. (N=3 tumors per clone).

There are many immune cell types in the tumor microenvironment that play critical anti-tumor and pro-tumor roles^25^. To achieve a more comprehensive assessment of immune cell infiltration in the untreated tumors, we used CIBERSORTx computational deconvolution of the bulk-RNAseq data to obtain absolute proportion scores of multiple immune cell types **(Fig. 5b)**. In general, C1 followed by C11, C22 and C23 showed the highest absolute immune infiltration. Consistent with the CD3 immunostaining, T cells were the most abundant population in C1 tumors, with an estimated proportion of CD8^+^ T cells significantly greater than tumors from other clonal sublines (except for C18). On the other hand, C11, C22 and C23 tumor immune infiltrates were predominantly myeloid cells and an elevated proportion of M2 macrophages was observed, uncovering a marked immunosuppressive TME in these clones^25^. In contrast, the tumors from C12 and C15 were estimated to have the least number of immune infiltrates with both anti- and pro-tumor functions, suggesting an immune-depleted or “cold” microenvironment. Overall, these results underscore a remarkable variety of TME compositions in the tumors from the clonal sublines.

To further characterize the microenvironment of the untreated tumors, we analyzed the expression of 29 immune and stroma related gene signatures recently compiled and utilized to predict melanoma patient survival and therapeutic response^26^ **(Fig. 5c-d)**. The expression of the 29 signatures in the tumors resulted in four branching groups, which we described as: (1) inflamed, (2) inflamed-T cell dysfunctional, (3) immune depleted-fibrotic; and (4) T cell exclusion-fibrotic. Tumors in the inflamed (1) category (e.g., C1, C8, C14) exhibited increased expression of genes related to anti-tumor immune infiltration (e.g., NK, B and T cells, MHC-I, Th1 and M1 signatures, co-stimulatory ligands/receptors, and antitumor cytokines) together with decreased expression of pro-tumor signatures (e.g., myeloid cells, tumor-associated macrophages [TAMs] and cancer-associated fibroblasts [CAFs], and protumor cytokines). Tumors in the inflamed-T cell dysfunctional (2) category (e.g., C11, C22, C23) presented upregulation of both anti-tumor (e.g., NK and T cells) and pro-tumor (e.g., Treg, TAMs and neutrophils) immune profiles. Immune depleted-fibrotic (3) tumors (e.g., C4, C12, C15) had increased expression of signatures associated with fibrosis (e.g., CAFs, matrix remodeling, endothelium), accompanied by decreased expression of immune infiltrate profiles (e.g., effector cells, Treg and TAMs). Tumors in the T cell exclusion-fibrotic (4) category (e.g., C8, C11, C22) also had increased expression of fibrosis and pro-tumor immune cell signatures (e.g., matrix, CAFs, TAMs, Tregs, protumor cytokines and checkpoint molecules) along with low effector cells and antitumor cytokines profiles. Consistent with the results from CIBERSORTx and CD3 IHC, untreated tumors from clonal subline C1 presented the highest level of anti-tumor inflammation, whereas C11 presented immune exclusion and dysfunction phenotypes.

In relation to their previously described developmental states *in vivo*, the TMEs of undifferentiated clonal sublines (e.g., C11, C23) were predominantly characterized as T cell exclusion or dysfunctional, while the TMEs of more differentiated clonal sublines (e.g., C14, C18, C15) primarily demonstrated immune depleted-fibrotic phenotypes **(Fig. 5d)**. Additionally, the undifferentiated, T cell exclusion-fibrotic tumors (e.g., C11, C23) were associated with downregulation of ‘Apical junction’ and ‘Myogenesis’ Hallmark pathways related to cell migration, while the more differentiated, immune depleted-fibrotic tumors were associated with downregulation of pathways such as ‘E2F targets’ and ‘G2M checkpoint’ related to proliferation. Transitory, inflamed tumors (e.g., C1) exhibited upregulation of proliferation pathways (e.g., ‘E2F targets’ and ‘G2M checkpoint’) and downregulation of pathways related to metastasis and immune responses (e.g., ‘Epithelial mesenchymal transition’, ‘Allograft rejection’). Altogether, these results illustrate the complex interplay between melanoma intrinsic phenotypic states and the immune and stromal populations of the TME.

### Diverse response of the selected clones to anti-CTLA-4 therapy

Based on our molecular characterization of the clones in culture and *in vivo*, we selected 4 sublines (C1, C11, C14 and C15) representing diverse genomic, transcriptomic, and immune profiles to test anti-CTLA-4 efficacy. We implanted 1 million cells of each subline into syngeneic C57BL/6 mice and when tumors reached an average of 75 mm^3^, mice were randomized and treated with four doses of anti-CTLA-4 or isotype antibody over the course of two weeks (**Fig. 6a**). Tumor size was measured until endpoint, which defined the survival time (see **Methods**). C1 tumors exhibited the greatest response to anti-CTLA-4, with 30% of tumors showing full regression or stable volume and 30% with over a 4-month growth delay, resulting in significantly improved overall survival. Mice implanted with C14 presented 40% response rate leading to significant improved survival compared to the isotype control group, but all tumors relapsed between 75 to 120 days. Anti-CTLA-4 had a minor effect on C15-implanted mice with only one tumor showing a 21-day growth delay. Finally, C11 tumors were fully resistant to anti-CTLA-4 treatment as no significant tumor growth delay or survival improvement were observed **(Fig. 6b-c)**. These results reveal the phenotypic diversity of the M4 clonal sublines in line with their molecular heterogeneity, which recapitulates the ITH and the variety of responses to ICB demonstrated by the M4 parental model.

**Fig. 6.**
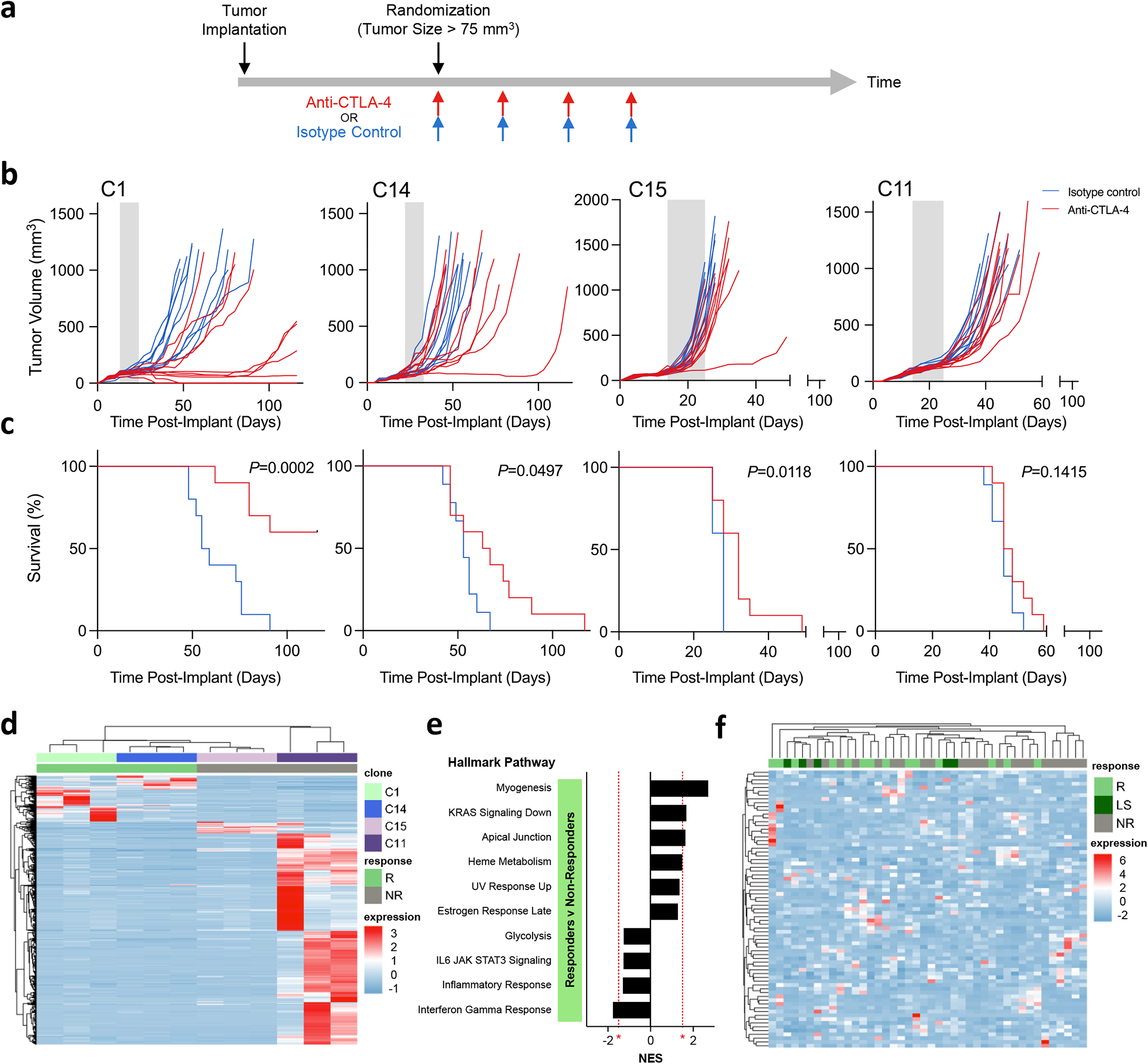
Diverse response of the clones to anti-CTLA-4 therapy. **a,** 1.0 x 10^6^ melanoma cells from clonal sublines C1, C14, C15, and C11 were implanted subcutaneously into immunocompetent syngeneic C57BL/6 mice. When tumors reached an average of 75 mm^3^, mice were randomized and treated with anti-CTLA-4 or isotype antibody twice per week. **b,** Tumor growth curves of the clones implanted in mice treated with anti-CTLA-4 (red lines, N=10) or isotype control (blue lines, N=10). Shaded gray area indicates treatment period. **c,** Kaplan-Meier survival curves from Fig.6a. Two-tailed *P* values from log-rank (Mantel-Cox) test are indicated. **d,** Heat map of differentially expressed genes (DEGs) between untreated responder (R) tumors (C1 and C14) versus non-responder (NR) tumors (C15 and C11) from DESeq2 analysis. Scale bar represents gene expression values after z-score normalization of FPKM. **e,** GSEA of Hallmark pathways comparing untreated responder versus non-responder tumors from the clones. Normalized enrichment score (NES) is shown, red asterisks and dashed lines indicate thresholds of significance. **f,** Heat map showing unsupervised hierarchical clustering of Van Allen data set^27^ (pre-treatment samples of metastatic melanoma patients treated with anti-CTLA-4) by their expression of the DEGs in Fig.6d. (R: responder, LS: long-survival, NR: non-responder). Scale bar represents gene expression values after z-score normalization of RPKM. Genes with RPKM expression values less than 10 in more than ∼10 percent of patients (N=4/42) were filtered out.

Next, we examined whether the ICB outcomes observed for C1, C14, C15 and C11 were related with their immunogenicity. For evaluation of *in vivo* anti-tumor T cell responses, we implanted the four clonal sublines into athymic NUDE mice **(Fig.S6a-b)**. A significant increase in tumor growth in the NUDE mice was observed for C1, C14, and C15 tumors, indicating the presence of antitumoral T cell responses in immunocompetent mice against these clonal sublines. There was no difference in tumor growth observed for C11 tumors in immunocompetent versus NUDE mice, explaining the resistance of C11 to anti-CTLA-4. Next, we evaluated the expression of neoantigens and antigen presentation genes in the tumors **(Fig.S6c-e)**. MHBR-I/II scores were calculated to analyze the predicted binding of the expressed neoantigens to MHC-I/II receptors **(Fig.S6c-d)**. C1 had the greatest number of predicted neoantigens binding to both MHC-I and MHC-II, which explains in part the antitumor activity observed in line with its response to anti-CTLA-4. The antigen-presentation pathway was found to be functional with no substantial differences in the expression of MHC-I/II related genes in untreated tumors from these clones **(Fig.S6e)**. Overall, these results demonstrate the correlation of tumor immunogenicity with ICB efficacy in the four clonal sublines, which is consistent with previous preclinical and clinical studies^13,27^.

### The transcriptomic profiles of the clonal sublines distinguish ipilimumab responder and non-responder melanoma patients

After observing the responses to ICB, we performed differential gene expression analysis to compare clones that responded to anti-CTLA-4 therapy (R: C1 and C14) versus non-responder sublines (NR: C15 and C11) **(Fig. 6d)**. GSEA using Hallmark pathways revealed that untreated tumors from sensitive clonal sublines had increased expression of genes associated with ‘Myogenesis’, ‘KRAS signaling down’, and ‘Apical junction’ pathways, while those from resistant clones had greater expression of genes related to ‘Interferon gamma response’ **(Fig. 6e)**. These results are consistent with recent reports demonstrating worst outcomes of patients receiving ICB therapy when melanomas presented chronic IFNγ pathway activation before treatment^7,28,29^. To validate our findings, we examined the expression of the differentially expressed genes from the clonal sublines in metastatic melanoma patients treated with ipilimumab (Van Allen dataset; **Fig. 6f**). Unsupervised hierarchical clustering based on the gene set from the clones resulted in 3 clusters, the first one enriched on responder (R) and long-survival (LS) patients, the second with mixed samples, and the third with only non-responder (NR) patients (**Fig. 6f**). These results highlight the utility of using M4 clonal sublines to understand clinical outcomes and identify improved treatment strategies for melanoma patients.

## DISCUSSION

The relevance of ITH for metastatic progression and therapeutic resistance is well established^6^. While metastatic colonization is thought to result from the expansion of aggressive clones, it is unclear whether cancer treatment leads to the selection of drug-resistant clones or the adaptation of pooled drug-tolerant cells that refuel ITH of relapsed tumors^5,8,10^. Similarly, the relative impact of genomic alterations versus non-genetic factors, such as transcriptional plasticity, on the therapeutic response remains unknown. Moreover, these factors are most likely context dependent in relation to cancer type and stage or specific drug treatment, and their dissection at the single cell level is critical for better clinical management of tumor relapse and personalized medicine. We have generated a unique experimental system to study these determinants consisting of 24 single cell-derived, low passage sublines from the M4 GEM model of cutaneous melanoma. M4 reliably mimics melanoma etiology and behavior and is well suited for investigating intra-tumoral dynamics due to its genetic and phenotypic heterogeneity, including mixed response to ICB treatment. Importantly, these sublines exhibit diverse and distinct molecular features (such as mutational landscapes, transcriptomic profiles and developmental states) and phenotypes (e.g., cell growth kinetics, immunogenicity and tumor formation potential, TME composition and ICB response).

Preclinical modeling, together with the most recent advances in single cell sequencing, genetic editing technologies, and computational analysis allows for the unraveling of cancer evolution complexity at multiple levels. Fully immunocompetent cancer models are particularly useful for conducting controlled preclinical studies on how cancer cells evolve during immunotherapy, which is not possible using clinical samples and data sets. However, due to experimental design and inter-species differences, most of the available preclinical models do not fully recapitulate human heterogeneity, especially at the genetic level^30–32^. To imitate the main etiological risk factor for melanoma initiation in humans, M4 was developed by exposing neonatal mice to relevant UV radiation^14^. This, together with the lack of strong oncogenic drivers, led to long latency and cancer evolution mirroring that of human melanomas. This approach ultimately resulted in a melanoma model with high TMB and ITH^13^.

The clonal sublines isolated from M4 exhibited genetic homogeneity but were highly diverse and distinct from one another, as demonstrated by phylogenetic analyses. Importantly, this confirmed the suitability of our experimental system for investigating genetic and non-genetic factors of melanoma growth and evolution. When cell proliferation and tumor growth in syngeneic and immunocompromised mice were evaluated, we found that T cell responses played a significant role in impairing the tumorigenicity of the clonal sublines. Not surprisingly, we found markedly increased TMB in clones unable to form tumors in fully immunocompetent mice (e.g., C19) or highly sensitive to ICB (e.g., C1), consistent with clinical observations and the potential use of TMB as a prognostic and predictive biomarker for ICB^33,34^. However, our results uncovered the relevance of specific mutational profiles, rather than total mutation or neoantigen load, for the tumor forming ability of melanoma cells. These findings agree with previous studies that associated immune surveillance with neoantigen heterogeneity and presentation^35,36^. Accordingly, this panel of clonal sublines has proven to be a powerful tool to identify functionally relevant and/or immunogenic mutations^16^.

There is growing evidence linking melanoma developmental states with resistance to immunotherapy^13,37,38^. Previous bulk-RNAseq analysis revealed that M4 melanoma represents a transitional developmental state, between neural crest-like and differentiated melanocytic phenotypes, in line with similar states found in patients^15^. However, whether this reflects a transitory state or the mixture of multiple states within a given tumor was unknown. The expression of canonical markers of melanocytic development and MPS scores revealed the wide range of differentiation states of the clonal sublines, suggesting that multiple developmental states coexist at the cellular level in M4 melanomas. Intriguingly, while most of the cells were predominantly in undifferentiated to neural crest states when grown in culture the majority of the clones presented transition or differentiated states *in vivo*. This is consistent with studies of human data sets that found most melanomas tend to be differentiated but at the same time contain subpopulations of undifferentiated cells^4^. Moreover, a high degree of inter- and intratumor heterogeneity was observed with respect to their global transcriptomic profiles and developmental states. These results demonstrated the remarkable plasticity of the clonal sublines and underscore the functional implications of the developmental states in the adaptation to the microenvironment *in vivo*.

We found diverse responses of the sublines to anti-CTLA-4, offering a platform for the study of the determinants of immunotherapy response. Tumors characterized as immunologically “hot” had higher levels of T cell infiltration and corresponded with improved ICB efficacy, whereas “cold” tumors lacked T cell infiltration and were associated with ICB resistance^24,39^. However, there are many immune cell populations that influence therapeutic responses, and several additional TME categories have been identified that are associated with patient prognosis and ICB efficacy, such as T cell dysfunction and exclusion^12,26,40^. Therefore, it is fundamental to understand the complete immune composition of tumors to predict and understand outcomes. TME characterization of untreated tumors from the clonal sublines resulted in four main subtypes ranging from immune-inflamed (e.g., C1 and C14) to T cell exclusion or immune depleted environments (e.g., C11, C14 and C15). But when we tested the response of representative clonal sublines to anti-CTLA-4, clones described as having both immune inflamed and depleted TMEs in untreated conditions (e.g., C1 and C14) were associated with response. In contrast, clones described as having T cell dysfunctional or exclusion TMEs (e.g., C11) were non-responsive to anti-CTLA-4, suggesting that tumors with established immunosuppressive environments correlate more strongly with resistance.

In connection with the differentiation status of the untreated tumors, undifferentiated clonal sublines (e.g., C11, C23) were largely described as T cell exclusion or dysfunctional, while more differentiated clonal sublines (e.g., C14, C18, C15) were predominantly described as immune depleted and fibrotic. Transitory status clones (e.g., C1, C4, C22) formed tumors that fell into all four TME subtypes. Interestingly, some clonal sublines, such as C22, exhibited remarkable phenotypic variability *in vivo,* with tumors falling into different transcriptional clades and TME subtypes. This provides evidence of crosstalk between the melanoma cells and involvement of host immunity determining tumor evolution and outcomes.

There are several approaches to track clonal evolution and explore heterogeneity-driven resistance mechanisms. Lineage tracing methods have been widely used to uncover melanoma development and progression via tracing cell migration, proliferation, and differentiation *in vivo^42–44^*. However, these strategies require the introduction of exogenous elements into the cells that generate limitations on performing functional analyses under immunocompetent conditions^45^. Here, we have used single cell-derived sublines to study how genetically homogenous populations can become transcriptionally plastic along tumor growth and ICB treatment. This system provided evidence of the critical role of neoantigens to trigger anti-tumor immune responses, uncovered the contribution of melanocytic developmental programs on melanoma adaptation and remodeling of the TME, and has proven to be a powerful tool for the discovery of correlates of ICB response in humans. The combination of anti-CTLA-4 (ipilimumab) and anti-PD-1 (nivolumab and pembrolizumab) treatment has shown significantly improved patient survival compared to monotherapy in clinical trials^46^, necessitating further investigation into the determinants of response under different treatment plans. The clonal sublines will be a useful tool to probe underlying mechanisms of response to monotherapy and combination ICB therapies (e.g., CTLA-4, PD-1/L1, LAG-3, TIM-3/3L targets).

## METHODS

### Cell lines

B2905 (M4) mouse cell line^13^ and the clonal sublines were cultured in RPMI supplemented with 10% of FBS and 1% L-Glutamine. To generate single-cell derived sublines, exponentially growing M4 cells were sorted in a 96-well plate by fluorescence-activated cell sorting (FACS) using a FACSAriaIII cytometer. 24 single cells were successfully cultured and expanded to create early-passage sublines. Authentication of all cell lines was performed by whole exome sequencing, HGF transgene genotyping and Spectral karyotyping (SKY). Cell lines were confirmed as Mycoplasma negative using a MycoAlert Mycoplasma Detection Kit (Lonza, LT07-418).

### Cell growth kinetics

IncuCyte live cell analysis was used for real-time quantitative live-cell imaging *in vitro*. Cell growth measurements were assessed by seeding 1.5 x 10^4^ cells of each clone per well of 24 well plates in triplicates. Plates were then placed into an IncuCyte S3 or SX5 (Sartorious) at 37 °C, 5% CO2, and 95% relative humidity. 16 phase-contrast images per well were taken every three hours for a total of five days. The IncuCyte Zoom software calculated a % confluence as a Phase Object Confluence. Doubling times were determined using GraphPad Prism 8.4.3 software by applying nonlinear regression ‘Exponential growth with log(population)’ on the linear phase of the curve: Y(ln(confluence)) = lnY0 + k*X and doubling time = ln(2)/k Experiments were repeated at least 3 times for each clone.

### *In vivo* tumor growth and treatment

The mice used in the preclinical studies were 6-12 weeks old female C57BL/6NCr (C57BL/6) or Athymic NCr-nu/nu (NUDE) mice, supplied by Charles River facility in NCI-Frederick, with no selection on the weight. They were housed in AAALAC-accredited cage as a group of five, with *ad libitum* food and water, and a 12-hour light cycle. All mouse experiments were performed in accordance with Animal Study Protocols approved by the Animal Care and Use Committee, NCI, NIH. NCI is accredited by the Association for Assessment and Accreditation of Laboratory Animal Care and follows the Public Health Service Policy on the Care and Use of Laboratory Animals. All animals used in this research project were cared for and used humanely according to the following policies: the US Public Health Service Policy on Humane Care and Use of Animals (2015); the Guide for the Care and Use of Laboratory Animals (2011); and the US Government Principles for Utilization and Care of Vertebrate Animals Used in Testing, Research, and Training (1985).

For tumorigenicity studies, 1.0×10^6^ melanoma cells from each cell line were implanted subcutaneously (s.c.) into C57BL/6 or NUDE mice and tumor growth was monitored until endpoint. For anti-CTLA-4 efficacy studies, 1.0×10^6^ melanoma cells from each cell line were implanted s.c. into C57BL/6 mice. When the tumors reached an average of 75 mm^3^, mice were randomized and treated with mouse anti-CTLA-4 (BioXCell, BP0164, clone 9D9) or mouse IgG2b as isotype control (BioXcell, BP0086). Antibodies were administered intravenously (i.v.) at a final dose of 10mg/Kg twice per week for a total of 4 doses. For all studies, tumor size and body weight were measured 2 times per week and the endpoint was established as the occurrence of one of the following events: (1) tumor size reached 1000 mm^3^; (2) tumor became ulcerated; and/or (3) the mouse showed moribund status or sickness behavior.

### Whole-exome sequencing

DNA was extracted from M4 cell line and the 24 clonal sublines using PureLink™ Genomic DNA Mini Kit (Invitrogen, cat# K1820-00). Mouse genomic DNA concentration was measured using Quant-iT PicoGreen dsDNA Assay Kit. 200ng of DNA was sheared by Covaris Instrument (E210) to ∼150-200bp fragments. Shearing was done in a Covaris snap cap tube (microTUBE Covaris p/n 520045) with the following parameters: duty cycle, 10%; intensity, 5; cycle burst, 200; time, 360 seconds at 4°C. Sample quality assessment/size was validated by Bioanalyzer DNA-High Sensitivity Kit (Agilent Technologies, #5067-4626).

Agilent SureSelectXT Library Prep ILM Kit (Agilent Technologies, #G9611A) was used to prepare the library for each sheared mouse DNA sample. DNA fragment’s ends were repaired, followed by adenylation of the 3’ end and then ligation of paired-end adaptor. Adaptor-ligated libraries were then amplified (Pre-capture PCR amplification: 98°C 2 minutes, 10 cycles; 98°C 30 seconds; 65°C 30 seconds; 72°C 1 minute; then 72°C 10 minutes, 4°C hold) by Herculase II fusion enzyme (Agilent Technologies, #600679). After each step, DNA was purified by Ampure XP beads (Beckmann Coulter Genomics #A63882). DNA Lo Bind Tubes, 1.5mL PCR clean (Eppendorf # 022431021) or 96-well plates were used to process the samples.

Samples were analyzed by bioanalyzer using the DNA-1000 Kit (Agilent Technologies, #5067-1504). The concentration of each library was determined by integrating under the peak of approximately 225-275bp. Then, each gDNA library (∼750-1000ng) was hybridized with biotinylated mouse specific capture RNA baits (SureSelectXT Mouse All Exon, catalog **#**5190-4641, 16 reactions) in the presence of blocking oligonucleotides. Hybridization was done at 65°C for 16 hours using SureSelectXT kit reagents. Bait-target hybrids were captured by streptavidin-coated magnetic beads (Dynabeads MyOne Streptavidin T1, Life Technologies, #6560) for 30 minutes at room temperature. Then, after a series of washes to remove the non-specifically bound DNA, the captured library was eluted in nuclease free water and half volume was amplified with individual index (Post-capture PCR amplification: 98°C 2 minutes; 10 cycles, 98°C 30 seconds; 57°C 30 seconds; 72°C 1 minute; then 72°C 10 minutes, 4°C hold). The Bioanalyzer DNA-High Sensitivity Kit was used to validate the size of the libraries prior to sequencing.

### Single-nucleotide variant analysis

All analyses were carried out on NIH biowulf2 high performance computing environment. Statistical analysis was carried out in R environment. Fastq sequence reads were mapped to the mouse reference genome mm10 with BWA or Bowtie. Single nucleotide variants (SNV) were identified using samtools mpileup or GATK HaplotypeCaller. Mouse germline single nucleotide polymorphisms (SNPs) were filtered out using the Sanger Mouse Genomes Project database for variants identified from whole genome sequencing of 36 mouse strains (ftp://ftp-mouse.sanger.ac.uk/current_snps/mgp.v5.merged.snps_all.dbSNP142.vcf.gz). Variants with a Phred-scaled quality score of <30 were removed. Variants that are present in normal spleen samples (in-house collection) were also removed. Variants were annotated with Annovar software to identify non-synonymous mutations.

### Single cell RNA-sequencing

To isolate cells for single cell RNA-sequencing, cells from each of the 24 established sublines *in vitro* were sorted into 96-well plates using FACS. Following the Smart-seq2 protocol^17^, the cells were lysed and full-length cDNA was generated via reverse transcription. The cDNA was amplified and then purified using Beckman Coulter AMPure XP beads. The quality and concentration of the cDNA from single cells of each subline, 3 samples were analyzed with the Agilent High Sensitivity D5000 ScreenTape Assay and 24 samples were analyzed with Quant-iT PicoGreen dsDNA Assay Kit. Based on these results, 8 samples from each of the 24 sublines were selected for tagmentation using the Illumina Nextera XT DNA sample preparation kit (Clones 1-12: Plate A/Set 1, Clones 13-24: Plate B/Set 2). The samples were diluted individually to reach optimal input concentration (<1ng) prior to tagmentation. After amplification of adapter-ligated fragments, the libraries were purified using Beckman Coulter AMPure XP beads. The quality and concentration of the libraries were checked using Agilent 2100 Bioanalyzer High Sensitivity DNA Assays prior to pooling and sequencing. The NovaSeq 6000 S1 system was used to sequence a total of 192 single cell libraries from the 24 clones.

### Seurat analysis

Cell clustering analyses were performed using the Seurat package with proper resolutions (2000 variable features, sctransform normalization)^47,48^. Cell cycle scoring and regression was performed to mitigate the effects of cell cycle heterogeneity using the CellCycleScoring function and mouse cell cycle genes. Uniform manifold approximation and projection (UMAP) was used to display identified cell clusters. UMAP cell clusters were annotated based on uniquely expressed genes, evolutionary relationships, clonal sublines, and MPS scores. The DoHeatmap function was used to visualize differential gene expression.

### Trisicell somatic variant calling and phylogeny tree building

After trimming and quality control process, raw reads were mapped to the mouse genome (GRCm38, mm10) using STAR (v2.7.3a) and then removal of duplicated mapped reads, indel realignment, and base re-calibration were performed. The pipeline for mutation calling from bulk exome sequencing and single-cell RNA sequencing are described previously^16^. Trisicell (Triple-toolkit for single-cell intratumor heterogeneity inference), was utilized to build tumor phylogeny based on mutations. It comprises three distinct methods: Trisicell-Boost effectively boosts any available method (PhISCS-BnB in this study) for tumor progression tree inference, to improve its scalability and speed; Trisicell-PartF computes the robustness of any inferred clade/subtree in a parameter free manner (in contrast to bootstrapping); and Trisicell-Cons compares trees generated from different omics data types or infers a consensus tree from them. The process of using Trisicell to infer expressed mutations from full-length scRNAseq data, generate mutation trees, and analyze tumor progression history was described previously^16^.

### Bulk RNA-sequencing

RNA was extracted from untreated tumors. Between 100ng to 1ug of total RNA was used to capture mRNA with oligo-dT coated magnetic beads. The mRNA was fragmented, and then a random-primed cDNA synthesis was performed. The RNA was fragmented into small pieces and the cleaved RNA fragments were copied into first strand cDNA using reverse transcriptase and random primers, followed by second strand cDNA synthesis using DNA Polymerase I and RNase H. The resulting double-strand cDNA was used as the input to a standard Illumina library prep with end-repair, adapter ligation and PCR amplification being performed to obtain a sequencing ready library. The final purified product was then quantitated by qPCR before cluster generation and sequencing.

### Gene expression analysis

The sequence reads in fastq format were aligned to the mouse reference genome mm10 using STAR and RSEM to obtain gene expression as Transcript Per Million (TPM) and the Fragments Per Kilobase of transcript per Million (FPKM) mapped reads.

#### Gene set enrichment analysis (GSEA)

The gene expression counts of the RNA sequencing data from the cell line-derived allografts were analyzed by DESeq2 (v1.25.1.) to compare between models and then the results were further analyzed by Gene set enrichment analysis (GSEA)^67^. For each comparison between two models, we identify enriched “Hallmark” pathways and considered those significant ones with an FDR<0.05.

#### Melanocytic plasticity signature (MPS) analysis

To calculate MPS scores the expression of the 45 genes in FPKM was weighted by 1 for up-regulated genes and -1 for the down-regulated genes and added. MPS scores were converted to *z-*scores across samples in each data set.

#### CIBERSORTx

For the analysis of immune cell population abundances from bulk RNA sequencing data, we used CIBERSORTx tool. In particular, we analyzed the absolute cell fraction of T cells in untreated tumors from the 24 clones and parental M4. Gene names were converted to human equivalent to create the mixture file. Impute Cell Fractions analysis module was used on CIBERSORTx. The signature matrix file LM22 was used for deconvolution, generated by the CIBERSORT authors from 22 leukocyte subsets profiled on the HGU133A microarray platform. B-mode batch correction was enabled and quantile normalization was disabled, which is the recommended method when using bulk-RNA sequencing data with the LM22 signature. The deconvolution was run in absolute mode and performed using 500 permutations for significance analysis. Default settings were used for all other parameters.

#### Immune signatures analysis

For the analysis of the tumor microenvironment using 29 immune signatures, ssGSEA was performed to generate a score per sample for each signature. A semi-supervised heatmap was generated using R, and the tumors were classified into four categories.

### Immunohistochemistry

Formalin-fixed and paraffin-embedded 5μm sections were stained with hematoxylin and eosin (HE), DCT (Pep8h), SOX10 (clone EP268, Cell Marque, 383R-15) or CD3 (clone CD3-12, BioRad, MCA1477) antibodies using standard immunohistochemistry methods. Antigen retrieval was performed in a pressure cooker using target retrieval buffer pH6 (Dako, S1699). Protein Blocking reagent (Dako, X0909) and Bloxall blocking solution (Vector, SP-6000-100) were used to block nonspecific proteins and endogenous peroxidase and alkaline phosphatase activities. The antibodies were incubated overnight at 4°C except for SOX10, which was incubated for 30 minutes at room temperature, in a humidity chamber. Antibody detection was performed using Impress AP Reagent anti-rabbit Ig (Vector, MP-5401) or Impress AP Reagent anti-rat IgG (Vector, MP-5444) and ImPACT Vector Red (Vector, SK-5105). Slides were counterstained with Mayer’s Hematoxylin (Electron Microscopy Sciences, 26043-06) and GIEMSA (New Comer Supply, 1120A). Digital images of whole-tissue sections were visualized using the HALO Image Analysis Platform.

#### HALO Image Analysis

HALO Image Analysis Platform v3.3.2541.300 was used to perform quantitative analysis of CD3 immunostaining and develop a machine learning-based classifier for melanin levels in whole-tumor sections. HALO CytoNuclear v2.0.9 algorithm was used to quantify the number of cells stained positive for CD3 protein as well as their staining intensity. Random Forest machine learning was used to develop a classifier for the regions of the tumors based on melanin levels.

### Statistical analyses

For *in vivo* therapeutic studies, tumors were measured independently by an animal technician, and the size (V) was calculated as following: V = 0.5*L*W^2^, in which L is the longer diameter and W is the shorter diameter. The survival time of the mice was defined as the duration from tumor implantation to the occurrence of one of the following events: (1) the tumor reached 2000 mm^3^; (2) tumor became ulcerated; or (3) the mouse showed moribund status or sickness behavior. The Kaplan-Meier survival analysis was performed using GraphPad Prism software (v7.0.1), and significance was analyzed by the Log-rank (Mantel-Cox) test. For vaccination studies, tumor onset time was defined as the post-implantation time when the tumor was observed to be palpable independently by an animal technician.

Unless otherwise indicated, all comparisons between two groups were analyzed by Mann-Whitney test with 95% confidence level using GraphPad Prism software (v7.0.1). For phenotypic analysis of tumor immune infiltrate in the four models, Kruskal-Wallis test followed by correction of multiple comparisons by Dunn’s test was performed in GraphPad Prism software (v7.0.1). For survival analysis of Van Allen^4^ and Hugo-Riaz^9,13^ data sets, Kaplan-Meier and Log-rank (Mantel-Cox) test was performed in the GraphPad Prism software (v7.0.1). All *P*-values obtained were two-tailed.

### Contact for reagents and resource sharing

Further information and requests for resources and reagents should be directed to G.M. Requests for the reagents developed in this study will require Material Transfer Agreement per NIH guideline.

## DATA AVAILABILITY

Whole exome sequencing, bulk and single-cell RNA sequencing raw data from this study will be deposited on Gene Expression Omnibus (GEO) upon publication.

## CODE AVAILABILITY

The custom code used in this manuscript will be available in the Online Methods section.

### Disclosure of Potential Conflicts of Interest

The authors declare no competing interests.

## Acknowledgments

This research was supported by funds from the NIH Intramural Research Program, a CCR Excellence in Postdoctoral Research Transition Award to E.P-G. and a CCR FLEX Synergy Award from the Center for Cancer Research (CCR) at the National Cancer Institute (NCI, NIH). We thank the NCI LGI Flow Cytometry Core, CCR Single Cell Analysis Facility (SCAF), Jennifer Dwyer and Noemi Kedei.

## Notes

### Competing Interest Statement

The authors have declared no competing interest.

## References

1 Stratton, M. R., Campbell, P. J. & Futreal, P. A. The cancer genome. Nature 458, 719–724 (2009). https://doi.org:10.1038/nature07943

2 da Silva-Diz, V., Lorenzo-Sanz, L., Bernat-Peguera, A., Lopez-Cerda, M. & Munoz, P. Cancer cell plasticity: Impact on tumor progression and therapy response. Semin Cancer Biol 53, 48–58 (2018). https://doi.org:10.1016/j.semcancer.2018.08.009

3 Dagogo-Jack, I. & Shaw, A. T. Tumour heterogeneity and resistance to cancer therapies. Nat Rev Clin Oncol 15, 81–94 (2018). https://doi.org:10.1038/nrclinonc.2017.166

4 Tirosh, I. et al. Dissecting the multicellular ecosystem of metastatic melanoma by single-cell RNA-seq. Science 352, 189–196 (2016). https://doi.org:10.1126/science.aad0501

5 Shaffer, S. M. et al. Rare cell variability and drug-induced reprogramming as a mode of cancer drug resistance. Nature 546, 431–435 (2017). https://doi.org:10.1038/nature22794

6 Vitale, I., Shema, E., Loi, S. & Galluzzi, L. Intratumoral heterogeneity in cancer progression and response to immunotherapy. Nat Med 27, 212–224 (2021). https://doi.org:10.1038/s41591-021-01233-9

7 Biswas, A. & De, S. Drivers of dynamic intratumor heterogeneity and phenotypic plasticity. Am J Physiol Cell Physiol 320, C750–C760 (2021). https://doi.org:10.1152/ajpcell.00575.2020

8 McGranahan, N. & Swanton, C. Clonal Heterogeneity and Tumor Evolution: Past, Present, and the Future. Cell 168, 613–628 (2017). https://doi.org:10.1016/j.cell.2017.01.018

9 Riaz, N. et al. Tumor and Microenvironment Evolution during Immunotherapy with Nivolumab. Cell 171, 934–949 e916 (2017). https://doi.org:10.1016/j.cell.2017.09.028

10 Rambow, F. et al. Toward Minimal Residual Disease-Directed Therapy in Melanoma. Cell 174, 843–855 e819 (2018). https://doi.org:10.1016/j.cell.2018.06.025

11 Sharma, P., Hu-Lieskovan, S., Wargo, J. A. & Ribas, A. Primary, Adaptive, and Acquired Resistance to Cancer Immunotherapy. Cell 168, 707–723 (2017). https://doi.org:10.1016/j.cell.2017.01.017

12 Huang, A. C. & Zappasodi, R. A decade of checkpoint blockade immunotherapy in melanoma: understanding the molecular basis for immune sensitivity and resistance. Nat Immunol (2022). https://doi.org:10.1038/s41590-022-01141-1

13 Perez-Guijarro, E. et al. Multimodel preclinical platform predicts clinical response of melanoma to immunotherapy. Nat Med 26, 781–791 (2020). https://doi.org:10.1038/s41591-020-0818-3

14 Noonan, F. P. et al. Neonatal sunburn and melanoma in mice. Nature 413, 271–272 (2001).

15 Tsoi, J. et al. Multi-stage Differentiation Defines Melanoma Subtypes with Differential Vulnerability to Drug-Induced Iron-Dependent Oxidative Stress. Cancer Cell 33, 890–904 e895 (2018). https://doi.org:10.1016/j.ccell.2018.03.017

16 Mehrabadi, F. R. et al. Profiles of expressed mutations in single cells reveal patterns of tumor evolution and therapeutic impact of intratumor heterogeneity. bioRxiv, 2021.2003.2026.437185 (2021). https://doi.org:10.1101/2021.03.26.437185v2

17 Picelli, S. et al. Full-length RNA-seq from single cells using Smart-seq2. Nat Protoc 9, 171–181 (2014). https://doi.org:10.1038/nprot.2014.006

18 Torre, E. A. et al. Genetic screening for single-cell variability modulators driving therapy resistance. Nat Genet 53, 76–85 (2021). https://doi.org:10.1038/s41588-020-00749-z

19 Wolf, Y. et al. UVB-Induced Tumor Heterogeneity Diminishes Immune Response in Melanoma. Cell 179, 219–235 e221 (2019). https://doi.org:10.1016/j.cell.2019.08.032

20 Linnemann, C. et al. High-throughput epitope discovery reveals frequent recognition of neo-antigens by CD4+ T cells in human melanoma. Nat Med 21, 81–85 (2015). https://doi.org:10.1038/nm.3773

21 van Rooij, N. et al. Tumor exome analysis reveals neoantigen-specific T-cell reactivity in an ipilimumab-responsive melanoma. J Clin Oncol 31, e439–442 (2013). https://doi.org:10.1200/JCO.2012.47.7521

22 Marty, R. et al. MHC-I Genotype Restricts the Oncogenic Mutational Landscape. Cell 171, 1272–1283 e1215 (2017). https://doi.org:10.1016/j.cell.2017.09.050

23 Kim, Y. J. et al. Melanoma dedifferentiation induced by IFN-γ epigenetic remodeling in response to anti-PD-1 therapy. The Journal of clinical investigation 131, e145859 (2021). https://doi.org:10.1172/JCI145859

24 Farhood, B., Najafi, M. & Mortezaee, K. CD8(+) cytotoxic T lymphocytes in cancer immunotherapy: A review. J Cell Physiol 234, 8509–8521 (2019). https://doi.org:10.1002/jcp.27782

25 Lei, X. et al. Immune cells within the tumor microenvironment: Biological functions and roles in cancer immunotherapy. Cancer Lett 470, 126–133 (2020). https://doi.org:10.1016/j.canlet.2019.11.009

26 Bagaev, A. et al. Conserved pan-cancer microenvironment subtypes predict response to immunotherapy. Cancer Cell 39, 845–865 e847 (2021). https://doi.org:10.1016/j.ccell.2021.04.014

27 Van Allen, E. M. et al. Genomic correlates of response to CTLA-4 blockade in metastatic melanoma. Science 350, 207–211 (2015). https://doi.org:10.1126/science.aad0095

28 Benci, J. L. et al. Tumor Interferon Signaling Regulates a Multigenic Resistance Program to Immune Checkpoint Blockade. Cell 167, 1540–1554 e1512 (2016). https://doi.org:10.1016/j.cell.2016.11.022

29 Shin, D. S. et al. Primary Resistance to PD-1 Blockade Mediated by JAK1/2 Mutations. Cancer Discov 7, 188–201 (2017). https://doi.org:10.1158/2159-8290.CD-16-1223

30 Rebecca, V. W., Somasundaram, R. & Herlyn, M. Pre-clinical modeling of cutaneous melanoma. Nature Communications 11, 2858 (2020). https://doi.org:10.1038/s41467-020-15546-9

31 Day, C.-P., Merlino, G. & Van Dyke, T. Preclinical Mouse Cancer Models: A Maze of Opportunities and Challenges. Cell 163, 39–53 (2015). https://doi.org/10.1016/j.cell.2015.08.068

32 Patton, E. E. et al. Melanoma models for the next generation of therapies. Cancer Cell 39, 610–631 (2021). https://doi.org:10.1016/j.ccell.2021.01.011

33 Wu, Y. et al. The Predictive Value of Tumor Mutation Burden on Efficacy of Immune Checkpoint Inhibitors in Cancers: A Systematic Review and Meta-Analysis. Frontiers in Oncology 9 (2019). https://doi.org:10.3389/fonc.2019.01161

34 Klempner, S. J. et al. Tumor Mutational Burden as a Predictive Biomarker for Response to Immune Checkpoint Inhibitors: A Review of Current Evidence. The Oncologist 25, e147–e159 (2019). https://doi.org:10.1634/theoncologist.2019-0244

35 McGranahan, N. et al. Clonal neoantigens elicit T cell immunoreactivity and sensitivity to immune checkpoint blockade. Science 351, 1463–1469 (2016). https://doi.org:doi.10.1126/science.aaf1490

36 Łuksza, M. et al. A neoantigen fitness model predicts tumour response to checkpoint blockade immunotherapy. Nature 551, 517–520 (2017). https://doi.org:10.1038/nature24473

37 Ballotti, R., Cheli, Y. & Bertolotto, C. The complex relationship between MITF and the immune system: a Melanoma ImmunoTherapy (response) Factor? Mol Cancer 19, 170 (2020). https://doi.org:10.1186/s12943-020-01290-7

38 Liu, D. et al. Evolution of delayed resistance to immunotherapy in a melanoma responder. Nat Med 27, 985–992 (2021). https://doi.org:10.1038/s41591-021-01331-8

39 Kather, J. N. et al. Topography of cancer-associated immune cells in human solid tumors. eLife 7, e36967 (2018). https://doi.org:10.7554/eLife.36967

40 Jiang, P. et al. Signatures of T cell dysfunction and exclusion predict cancer immunotherapy response. Nat Med (2018). https://doi.org:10.1038/s41591-018-0136-1

41 Hamid, O. et al. A prospective phase II trial exploring the association between tumor microenvironment biomarkers and clinical activity of ipilimumab in advanced melanoma. J Transl Med 9, 204 (2011). https://doi.org:10.1186/1479-5876-9-204

42 Köhler, C. et al. Mouse Cutaneous Melanoma Induced by Mutant BRaf Arises from Expansion and Dedifferentiation of Mature Pigmented Melanocytes. Cell Stem Cell 21, 679–693.e676 (2017). https://doi.org:10.1016/j.stem.2017.08.003

43 Moon, H. et al. Melanocyte Stem Cell Activation and Translocation Initiate Cutaneous Melanoma in Response to UV Exposure. Cell Stem Cell 21, 665–678.e666 (2017). https://doi.org:10.1016/j.stem.2017.09.001

44 Sun, Q. et al. A novel mouse model demonstrates that oncogenic melanocyte stem cells engender melanoma resembling human disease. Nature Communications 10, 5023 (2019). https://doi.org:10.1038/s41467-019-12733-1

45 Kester, L. & van Oudenaarden, A. Single-Cell Transcriptomics Meets Lineage Tracing. Cell Stem Cell 23, 166–179 (2018). https://doi.org/10.1016/j.stem.2018.04.014

46 Rotte, A. Combination of CTLA-4 and PD-1 blockers for treatment of cancer. Journal of Experimental & Clinical Cancer Research 38, 255 (2019). https://doi.org:10.1186/s13046-019-1259-z

