## Supplemental figures for "Melanoma clonal subline analysis uncovers heterogeneity-driven immunotherapy resistance mechanisms"

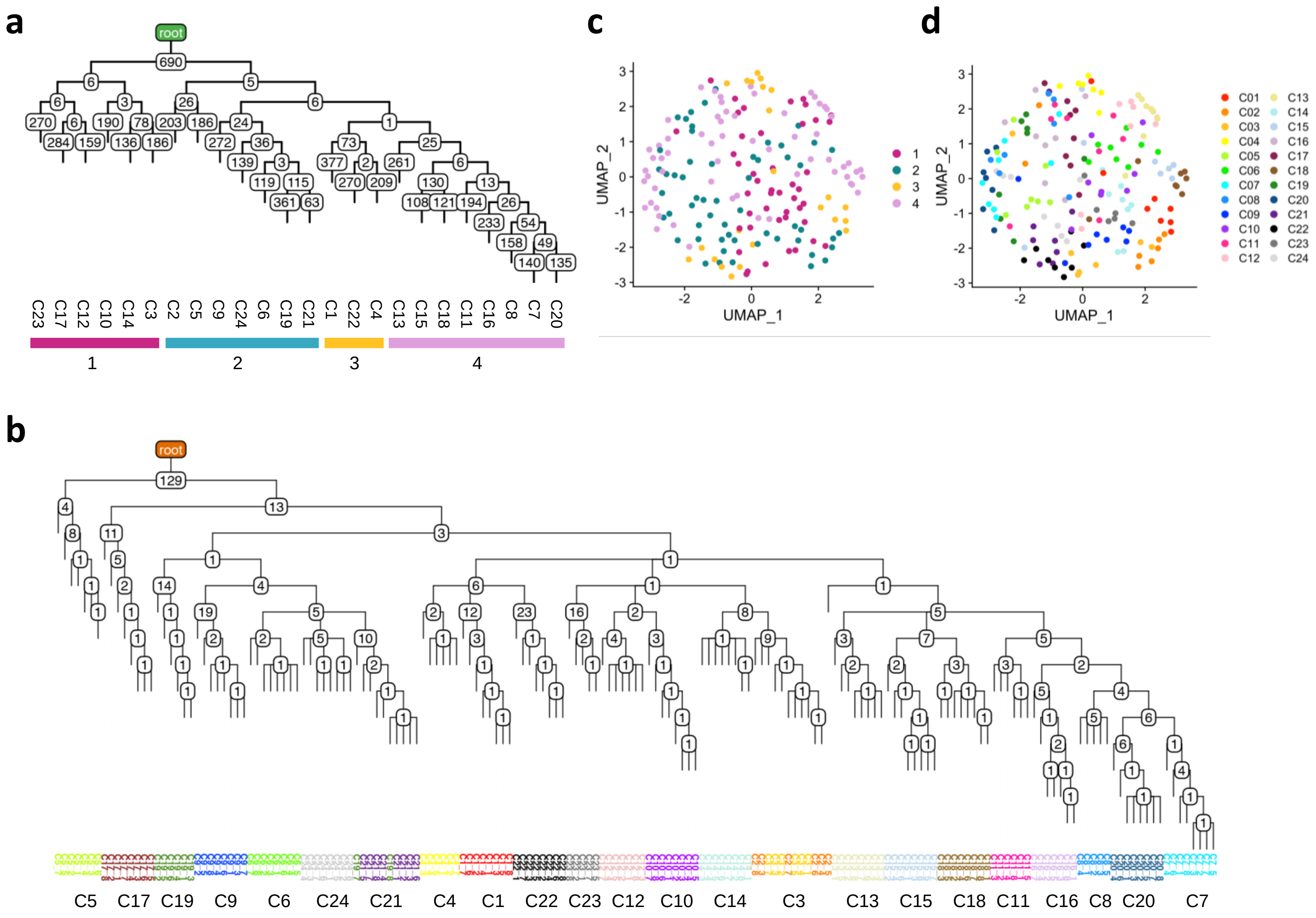


**Fig. S1 | Evolutionary relationships of the 24 clonal sublines inferred by *Trisicell* phylogenetic analysis. a,** Phylogenetic tree built by Trisicell-Boost using SNVs detected from whole exome sequencing (WES) data of the clones. The number of somatic mutations accumulated is shown for each node. The main branches of the tree are highlighted with color bars. **b,** Phylogenetic tree inferred by Trisicell-Boost using expressed SNVs called from scRNA-seq that were also detected on WES data (N=6-8 cells per clone). **c,** UMAP with cell color labels corresponding to the four branches of the WES-based phylogenetic tree from Fig.S1a. **d,** UMAP with cell color labels corresponding to the 24 clonal cell lines of the scRNA-seq-based phylogenetic tree from Fig S1b.

### Figure S2

Gruen et al.

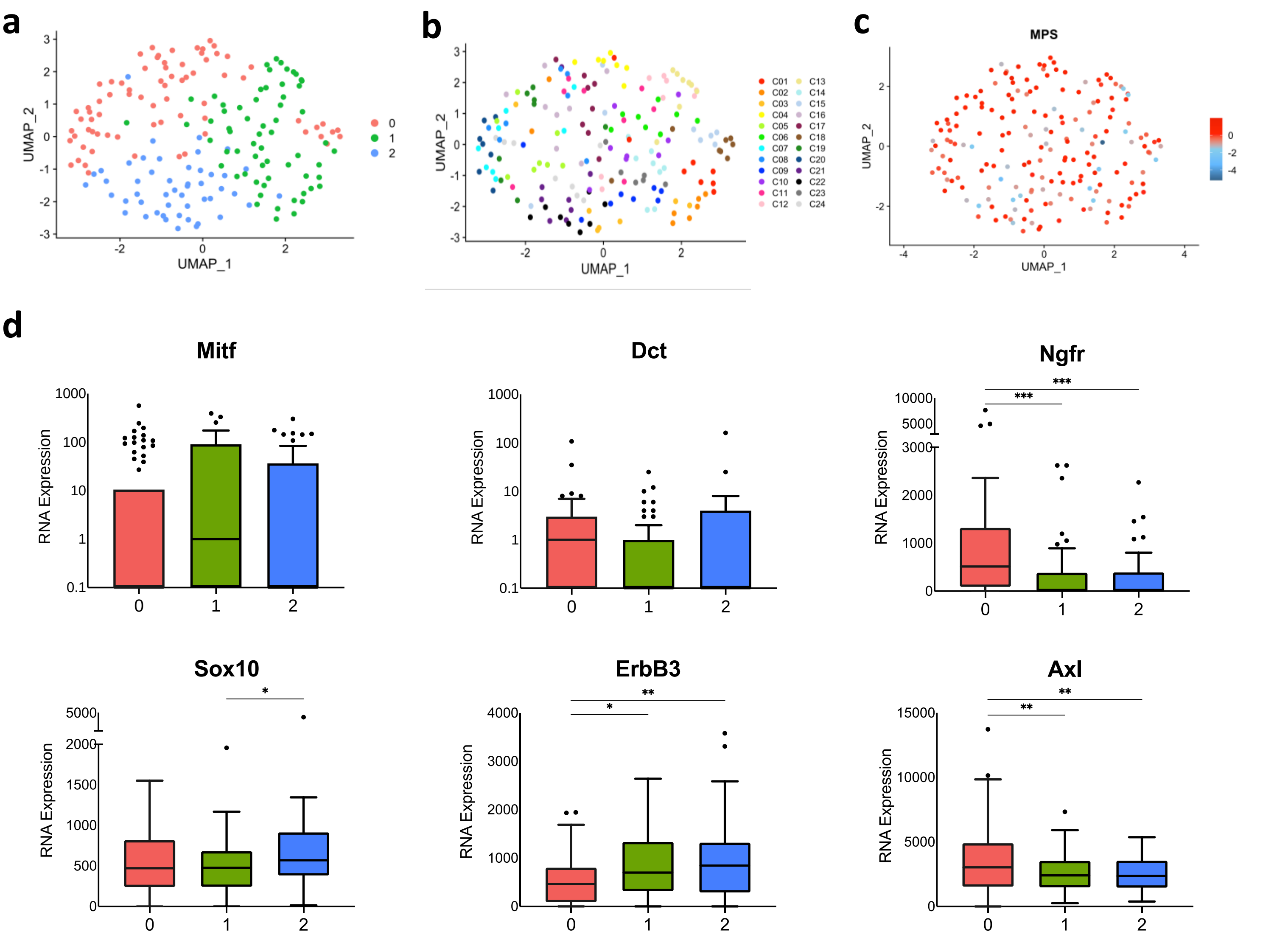


**Fig. S2 | Expression of melanocytic differentiation markers by the clonal sublines in culture. a,** UMAP with cell color labels corresponding to expression clusters (0-2). **b,** UMAP with cell color labels corresponding to the distinct clonal sublines (C1-C24). **c,** UMAP with cell color labels corresponding to Melanocytic Plasticity Signature (MPS) expression. Higher MPS scores represent undifferentiation, multipotency and/or neural crest stem cell properties, and lower MPS scores represent later stages of melanocytic differentiation. Scale bar represents z-score calculated with TPM normalized gene expression values. **d,** Box plots representing expression of canonical melanocytic marker genes (*Mitf, Dct, Ngfr, Sox10, ErbB3,* and *Axl*). RNA expression represented with SCTransform normalized gene expression values. Statistics shown for one-way ANOVA test adjusted for multiple comparisons (* indicates P≤0.05 , ** indicates P≤0.01, *** indicates P≤0.001).

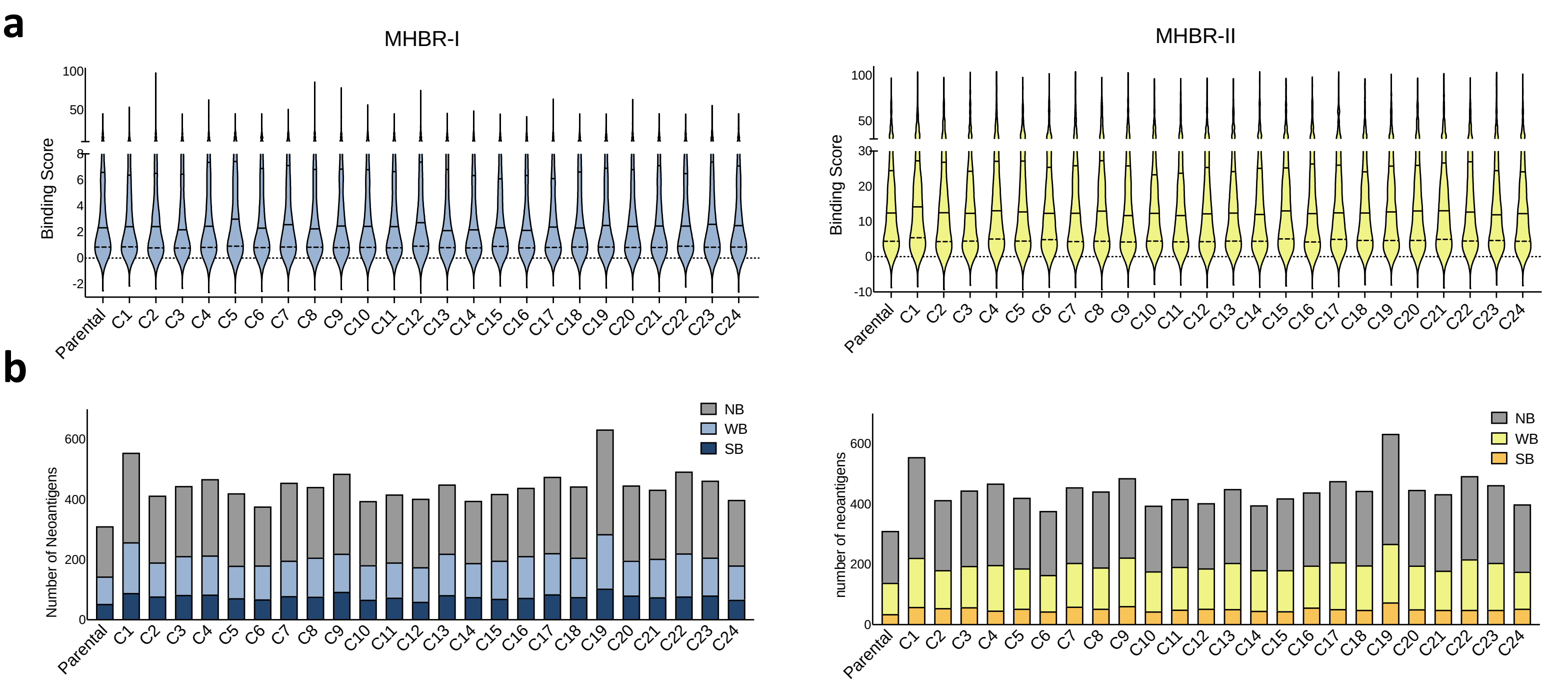


**Fig. S3 | MHC binding scores predict immunogenicity of cell line mutations. a,** Violin plots showing the distribution of MHBR-I (left) and MHBR-II (right) binding prediction scores of mutated genes from WES data of parental M4 and the 24 clonal sublines. Median values are indicated by solid black line and first and third quartiles marked by dashed black lines. **b,** Number of mutations predicted to bind MHBR-I (left) and MHBR-II (right) according to the following cutoffs: Strong-binding (SB)=0-0.5 for MHBR-I and 0-2 for MHBR-II; Weak-binding (WB)=0.5-2.0 for MHBR-I and 2-10 for MHBR-II; No binding (NB)=2.0-100 for MHBR-I and 10-100 for MHBR-II.

**a**

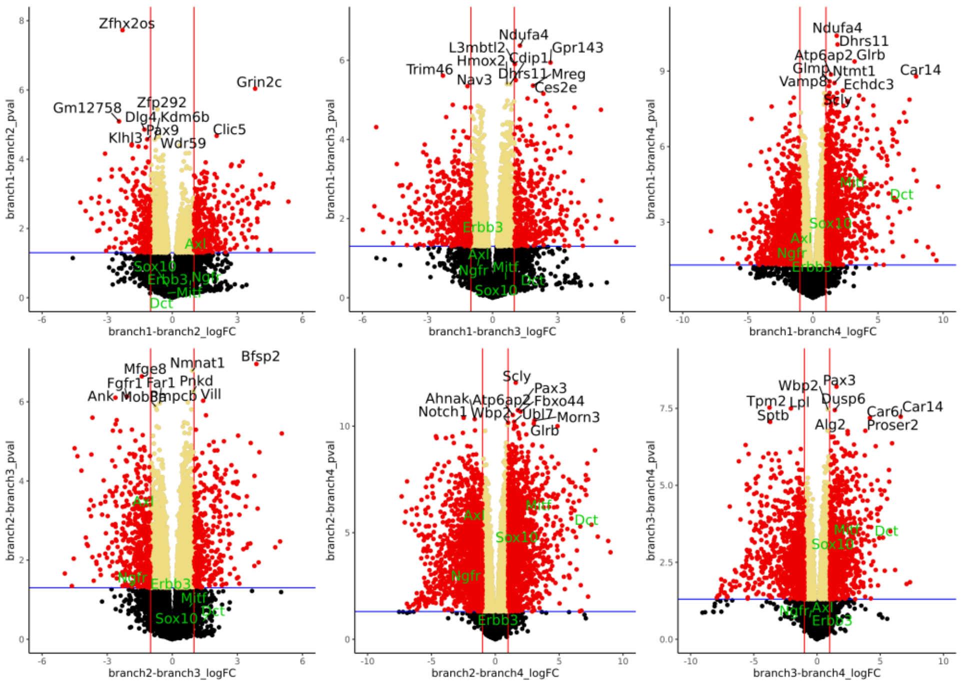


**b**

| I | II | III | IV |
|---|---|---|---|
| C4 | C18 | C16 | C8 |
| C1 | C22 | C8 | C22 |
| C1 | C14 | C22 | C23 |
| C1 | C12 | C4 | C23 |
| C14 | C12 | C12 | C8 |
| C14 | C15 | C16 | C23 |
| | C15 | C16 | C11 |
| | C15 | | C11 |
| | C4 | | C11 |
| | C18 | | |
| | C18 | | |

**Fig. S4 | Differential expression analysis from untreated tumors of the clones. a,** Volcano plots of differentially expressed genes (DEG) between branches (I-IV) from the heat map in Fig.4c. Top 10 DEG for each comparison indicated in black and differentiation marker genes (*Mitf, Dct, Ngfr, Sox10, Erbb3, Axl*) labeled in green. Blue and red lines represent designated thresholds (blue: p-value<0.05; red: logFC>1). **b,** Key indicating which tumors belong to each branch (I-IV).

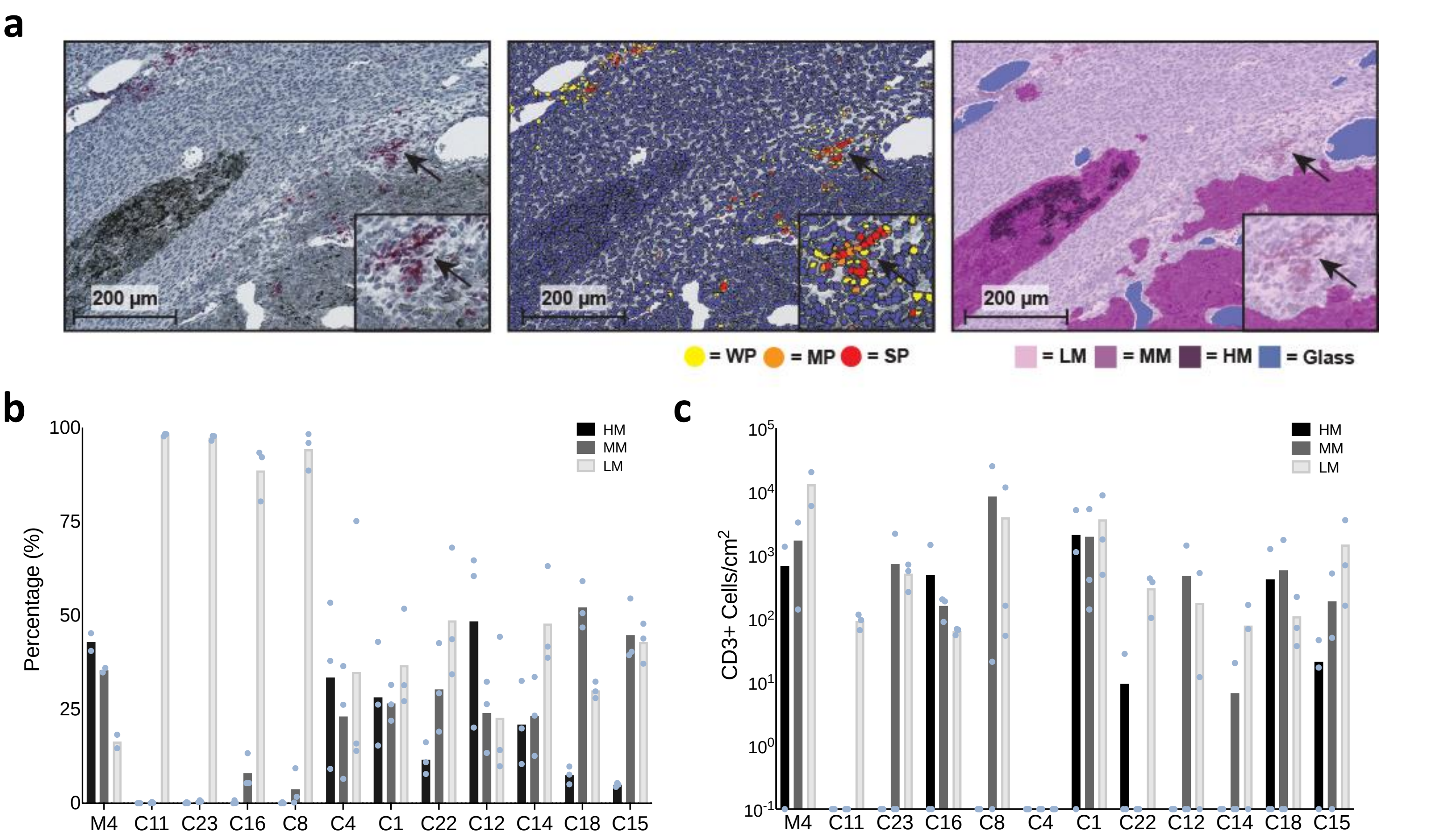


**Fig. S5 | Development of machine learning classifiers for the analysis of intratumoral T cells.** HALO Image Analysis Platform was used to perform quantitative analysis of CD3 immunostaining and develop a machine learning-based classifier for melanin levels in tumor sections. **a.** Representative images of the HALO classifiers. Left: CD3 immunostaining, arrows point $CD3^+$ cells (pink). Center: quantification of CD3 staining intensity and number of positive cells by HALO CytoNuclear algorithm. (WP: weak positive, MP: moderate positive, SP: strong positive, blue: CD3 negative cells). Right: classification of tumor areas based on the melanin content by Random Forest machine learning algorithm. (LM: low/no melanin, MM: moderate melanin, HM: high melanin). **b**. Quantification of melanin area in whole-tissue sections from the clones. **c**. Quantification of $CD3^+$ cells in each type of melanin area per clone. (N=3 tumors per clone and N=2 tumors for parental M4).

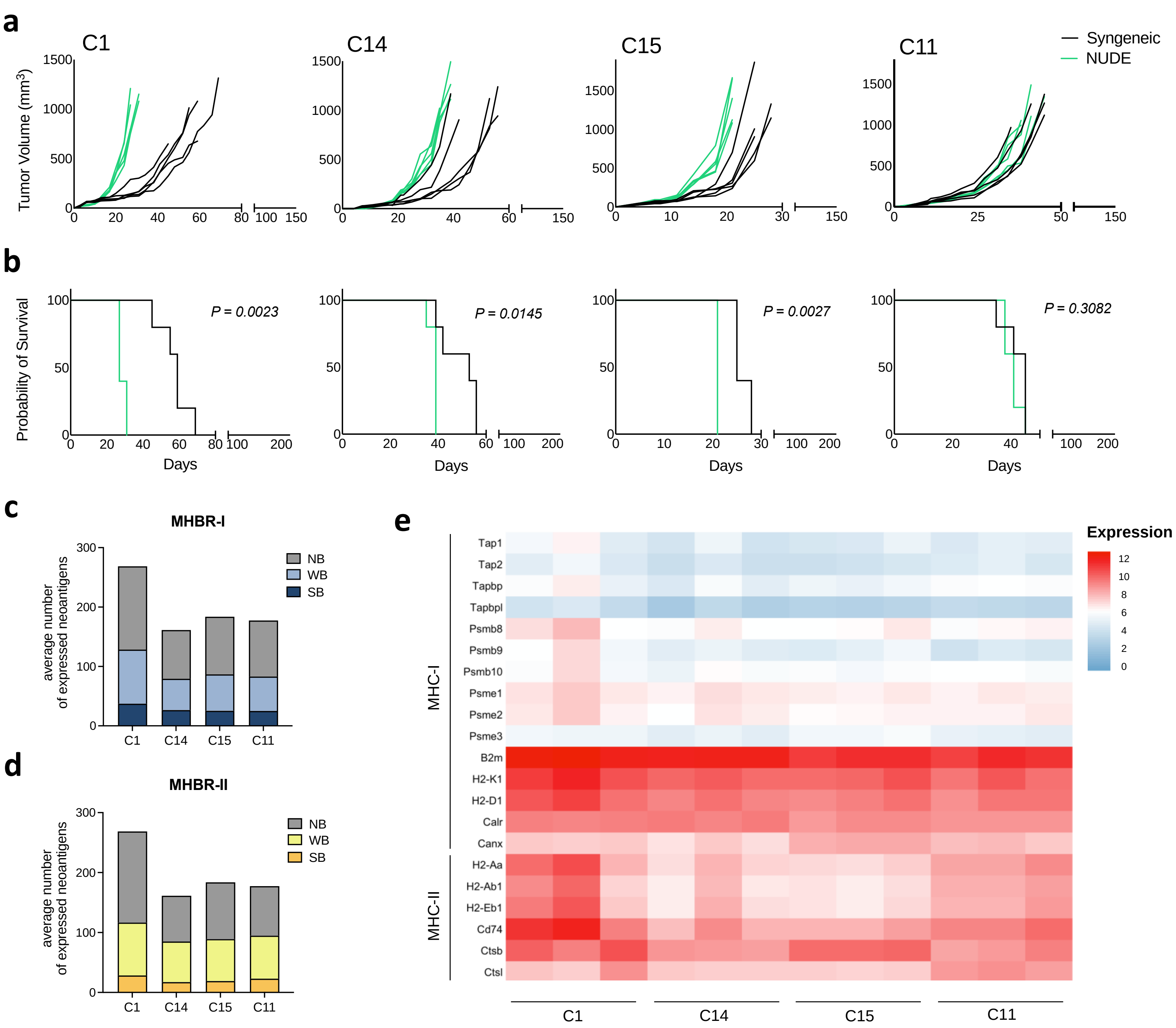


**Fig. S6 | Immunogenicity of tumors from clonal sublines C1, C14, C15, and C11. a,** 1.0 x $10^6$ melanoma cells from each clonal sublines were implanted subcutaneously into immunocompetent syngeneic C57BL/6 (black lines) or immunodeficient nude (green lines) mice. (N=5 mice per group). **b,** Kaplan-Meier survival curves from Fig.S6a. Two-tailed *P* values from log-rank (Mantel-Cox) test are indicated. **c-d,** MHBR-I and MHBR-II binding prediction of expressed mutated genes in the tumors from each clone (N=3 tumors per clone). Binding scores were calculated individually per tumor and results are shown for the average number of expressed mutant genes per clone falling into three categories (NB: non-binding, WB: weak-binding, SB: strong-binding). **e,** Expression of MHC class-I and class-II- related genes. Heat map depicts $Log_2$(FPKM+1) obtained from bulk RNA-seq of untreated tumors (N=3 per clone) .

#### Supplementary Tables provided upon request:

**Table S1. Nonsynonymous single nucleotide variants (SNVs) obtained from whole-exome sequencing of the 24 single-cell derived sublines.**

**Table S2.Differentially expressed genes between UMAP clusters related to Fig.1d.**

**Table S3. Gene Set Enrichment Analysis (GSEA) of differentially expressed genes between UMAP clusters related to Fig.1d.**

**Table S4. Differentially expressed genes between the tumors of each clade from the heatmap in Fig.4c.**
